# A Human Multi-Lineage Hepatic Organoid Model for Liver Fibrosis

**DOI:** 10.1101/2020.09.01.278473

**Authors:** Yuan Guan, Annika Enejder, Meiyue Wang, Zhuoqing Fang, Lu Cui, Shih-Yu Chen, Jingxiao Wang, Yalun Tan, Manhong Wu, Xinyu Chen, Patrik K. Johansson, Issra Osman, Koshi Kunimoto, Pierre Russo, Sarah C. Heilshorn, Gary Peltz

## Abstract

**Background:** To characterize fibrogenic mechanisms, genome engineering and a human hepatic organoid system was used to produce an *in vitro* model for human liver fibrosis.

**Methods and results:** Human hepatic organoids that were engineered to express the most common causative mutation for Autosomal Recessive Polycystic Kidney Disease (ARPKD) developed the key features of ARPKD liver pathology (abnormal bile ducts and hepatic fibrosis) in only 21 days. Second harmonic generation microscopy confirmed that the ARPKD mutation increased collagen abundance and thick collagen fiber production in hepatic organoids; and we demonstrated that these changes mirrored that occurring in ARPKD liver tissue. Transcriptomic and other analyses indicated that the ARPKD mutation generates cholangiocytes with increased TGFβ-associated pathway activation, which are actively involved in collagen fiber generation. The abnormal cholangiocytes promote the expansion of collagen-producing myofibroblasts with markedly increased PDGFRβ protein expression and an activated STAT3 signaling pathway. Moreover, the transcriptome of ARPKD organoid myofibroblasts resembled that of myofibroblasts in liver tissue obtained from patients with commonly occurring acquired forms of liver fibrosis. The involvement of the PDGFRB pathway was confirmed by the anti-fibrotic effect observed when ARPKD organoids were treated with PDGFRB inhibitors.

**Conclusions:** Besides providing mechanistic insight into the pathogenesis of congenital (and possibly acquired) forms of liver fibrosis, ARPKD organoids could also be used to test the anti-fibrotic efficacy of potential anti-fibrotic therapies.

## Introduction

Liver fibrosis is a pathological condition that results from extracellular matrix (**ECM**) accumulation in response to chronic liver injury ^1, 2^. Since excess ECM deposition eventually leads to loss of liver parenchymal cells and reduced liver function, fibrosis has severe and sometimes fatal complications. Although it is most commonly an acquired condition caused by viral infection or chronic alcohol exposure ^1, 3^, a few genetic diseases can cause liver fibrosis ^4^. While the rate of progression and histological features can vary in response to the different acquired or congenital causes, excess production of an altered ECM underlies all forms of liver fibrosis. This fibrotic state results from an interaction between parenchymal and nonparenchymal liver cells, and possibly involves infiltrating immune cells ^5–7^. The key non-parenchymal cell is the hepatocyte stellate cell (**HSC**), which is activated by a fibrogenic stimulus to transdifferentiate into a myofibroblast that is characterized by increased expression of a-smooth muscle actin (SMA), desmin (DES), and type I collagen (COL1A1) ^7–11^. Under normal conditions, the liver ECM consists of laminins, collagen (types I, III, and IV), and various proteoglycans ^12^; which provide important signals that maintain homeostatic conditions for liver cells. However, because myofibroblasts increase their production of fibril-forming collagen types I and III, collagen fibers become the most abundant component in the altered ECM of a fibrotic liver ^13, 14^. Thus, activated myofibroblasts and the collagens they produce are essential mediators of liver fibrogenesis. Irrespective of whether liver fibrosis has an acquired or congenital cause, no available treatments can prevent or reverse its progression if the underlying cause cannot be treated.

Autosomal Recessive Polycystic Kidney Disease (**ARPKD**: MIM263200) is a monogenic disorder (1 per 20,000 births) that primarily causes kidney and liver pathology ^15, 16^. The kidney disease is characterized by massive renal enlargement and collecting duct dilatation that progresses to renal failure and perinatal death in 30% of affected individuals ^17^. Despite the naming of this disease, liver disease is an important component of ARPKD. For the 70% that survive the perinatal period, liver disease becomes progressively more severe with age and becomes the major cause of morbidity and mortality ^15^. ARPKD liver disease is characterized by dilated intrahepatic bile ducts and a biliary fibrosis that is referred to as congenital hepatic fibrosis (**CHF**) ^16^. The lobular architecture is preserved in ARPKD liver disease ^18^. ARPKD is one of an expanding list of genetic disorders (ciliopathies) caused by dysfunction of primary cilia ^19^. It is caused by mutations within *polycystic kidney and hepatic disease-1* (*PKHD1*), which encodes a 4,074 amino acid multi-domain transmembrane protein (fibrocystin/polyductin, **FPC**) that is expressed in the primary cilia of renal tubular epithelial cells and cholangiocytes ^20, 21^. The vast majority of ARPKD subjects are compound heterozygotes with *PKHD1* mutations. Of the >800 *PKHD1* mutations that have been identified ^22–26^, the most common causative mutation is *Thr36Met* in exon 3; which accounts for 20% of all mutated alleles ^27^, and frequently appears in unrelated families of different ethnic origins ^22^. FPC is part of a protein complex ^28, 29^ that plays an important role as a chemosensor and mechanotransducer of extracellular environmental signals ^30–32^. Hence, *PKHD1* mutations cause cholangiocytes to have functionally abnormal primary cilia.

We present an *in vitro* model for a human liver fibrosis. To do this, we use our previously developed *in vitro* model system, where induced pluripotent stem cells (**iPSCs**) differentiate into human hepatic organoids (**HOs**) through stages that resemble those occurring during embryonic liver development ^33^. We demonstrate that this organoid system, when combined with genome editing technologies, reproduces ARPKD liver pathology. The ARPKD mutation causes biliary abnormalities and extensive fibrosis to develop in HOs in only 21 days. The ARPKD organoids have abnormal cholangiocytes and an expanded population of activated collagen-producing myofibroblasts. Moreover, transcriptomic similarities between the myofibroblasts in ARPKD organoids and those in liver tissue obtained from patients with commonly occurring acquired forms of liver fibrosis, suggests that this genetic program engenders changes that produce abnormalities that underly congenital and acquired forms of liver fibrosis.

*The materials and methods are provided in the supplement*.

## Results

### Characterization of a multi-lineage hepatic organoid

iPSCs can be induced to differentiate into hepatoblasts, which then differentiate into HOs in response to the sequential application of specific growth factor combinations that are added to the culture media (**Fig. S1a-b**). The HOs have sheets of hepatocytes, as well as cholangiocytes that are organized into epithelia around the lumina of bile duct-like structures, and these cell types can mediate many liver functions ^33^. We further confirm that HOs have primary cilium (**Fig. S1c**), which is an essential structure for analyzing genetic diseases that affect this organelle; and we demonstrate that organoids can synthesize pro-collagen and have the enzymatic machinery required for cross-linking collagen into thick fibers (**Fig. S1d**).

Single cell RNA sequencing (scRNA-Seq) has been a powerful method for characterizing cells within tissues ^34–37^, including liver ^38^. Therefore, scRNA-Seq analysis was performed on iPSC, hepatoblast, and hepatic organoid cultures. *As described in the supplemental note*, the cells at these differentiation stages could be separated into distinct clusters (**Figs. S2a-b**). As expected, the organoids had hepatocytes, cholangiocytes and bi-potential progenitor cells that can give rise to hepatocytes and cholangiocytes (**Figs. S2c**). Moreover, scRNA-Seq (**Figs. S2d-f**) and high-dimensional time of flight mass cytometry (CyToF) (**Figs. S2g-h**) data indicated that the organoids also contained other cell types including those that have resemble macrophages, endothelial cells and fibroblasts. The presence of these types of cells in organoids was confirmed by immunostaining (**Figs. S3a-c**). These results indicate that the HOs contain a more complex array of cells of different lineages, which form more complex structures than was noted in our prior studies ^33, 39^.

### An organoid model for ARPKD liver pathology

*PKHD1* mutations that cause amino acid substitutions are generally associated with a non-lethal presentation of ARPKD, while neonatal death tends to be associated with frame shift ^40^ or splice variant ^41^ alleles. Consistent with these clinical observations, we could not produce an iPSC line with a CRISPR engineered homozygous Ashkenazi founder mutation that induces a frame shift mutation (c.3761_3762delCCinsG) in *PKHD1* ^41^. However, we successfully engineered homozygous *PKHD^M36^* mutations into the genome of three different iPSC lines (C1-C3), each of which was produced from a different control individual (**Figs. 1a-b**). Inter-individual variation among donors is responsible for a large percentage of the phenotypic differences that have been observed in different iPSC lines ^42^. However, phenotypic differences that commonly occur in lines with the ARPKD mutation (but not in isogenic control lines) can be un-equivocally ascribed to the mutation (**Fig. 1c**). The morphology of HOs prepared from all three *PKHD^M36^* iPSC lines (which will be referred to as ARPKD lines) was altered in a characteristic manner relative to their isogenic controls (**Figs. 1d-e**). ARPKD organoids have an increased number of irregular bile ducts: bile duct structures occupied 30-40% of the area in ARPKD organoids versus 10-15% in control HOs. ARPKD organoids also had a markedly increased amount of ECM, which occupied 25~30% of the area in ARPKD HOs versus 0.3-0.5% of control HOs (**Figs. 1f, g**). The ECM increase was confirmed by immunostaining, which showed that an increased amount of collagen 1A (COL1A) was diffusely deposited in ARPKD organoids (**Fig. 1h**). Also, in contrast to the simple columnar morphology of the ductal epithelium in control organoids, ARPKD organoids had a disorganized ductal epithelium (**Fig. 1i-j**).

**Figure 1.**
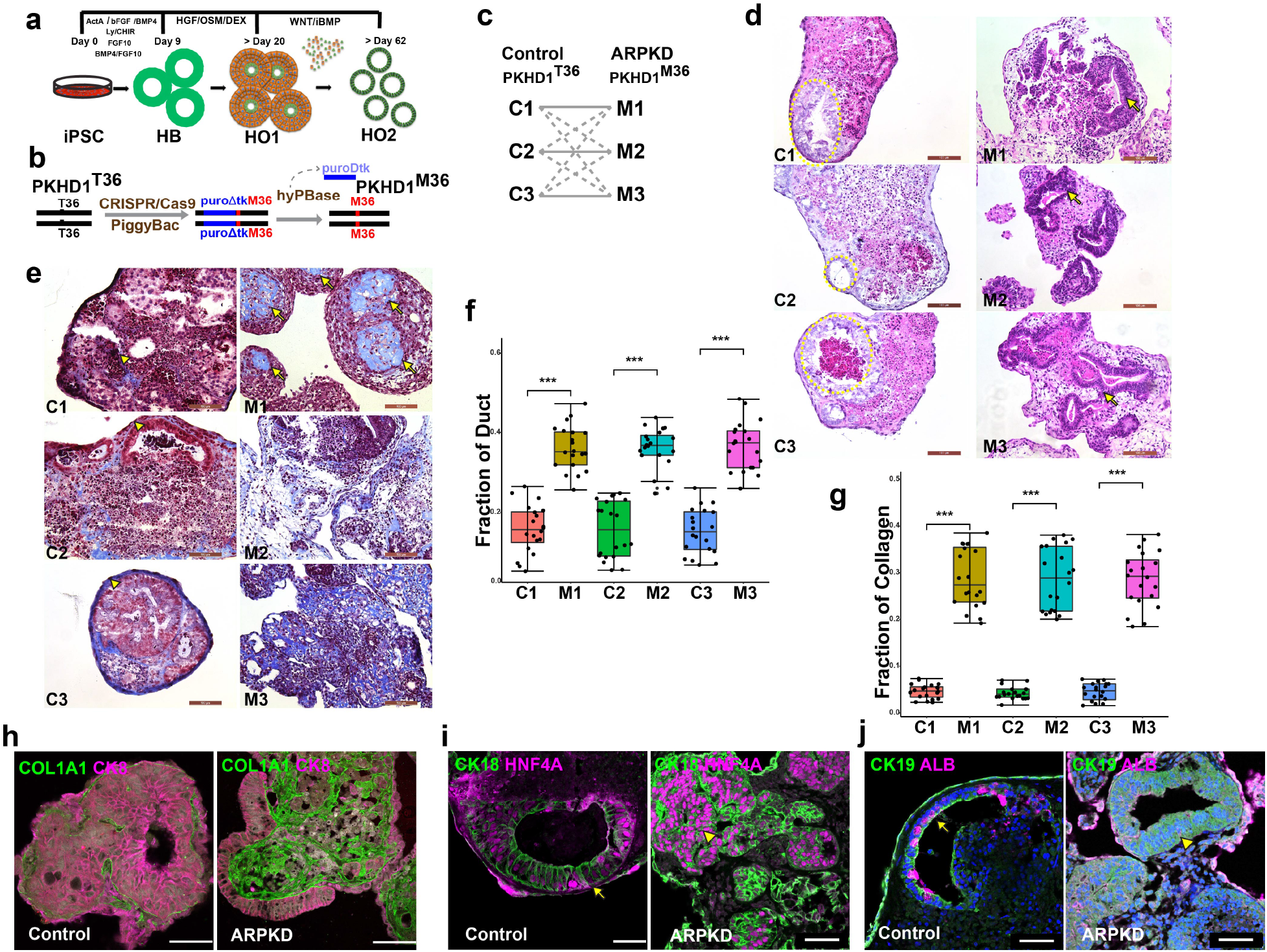
ARPKD organoids develop characteristic features of ARPKD liver disease. **a,** A schematic representation of the *in vitro* culture system and growth factors used to direct the differentiation of iPSCs into HOs. **b,** A diagram of the CRISPR engineering process used to produce iPSC lines with a homozygous ARPKD mutation (*PKHD1^36Met^*). **c,** Three iPSC lines with a homozygous *PKHD1^36Met^* mutation were produced from three unrelated control donors (C1-C3). This experimental design ensured that if a phenotypic difference was observed in a *PKHD1^36Met^* iPSC line relative to its isogenic control (solid lines), especially if it occurred in cells with different genetic backgrounds, the mutation could be unequivocally identified as the cause of the phenotypic difference. If this mutation had different effects in different iPSC lines, comparisons between the different lines (dotted lines) could identify the background differences that modified the effect of the mutation. **d, e** HOs prepared from the indicated isogeneic control (C1-3) or corresponding *ARPKD* (M1-3) iPSCs were stained with H&E (d) or the Trichrome stain (e). The trichrome stain shows the marked increase in collagen-rich connective tissue (blue regions, indicated by arrows) that appeared throughout all *ARPKD* HOs. In contrast, isogenic control organoids had a thin layer of connective tissue (indicated by arrowheads), which was often located at the periphery of the organoid. The dotted circles surround bile ducts. Scale bars are 100 μm. **f, g** The fraction of the total area within isogenic control and ARPKD HOs generated from the indicated donor, which was occupied by bile ducts (H&E stain) or by collagen. **h-j**, Control and ARPKD organoids were immunostained with antibodies to Collagen 1A (COL1A) and CK8 (**h**); or CK18, CK19, HNF4A and albumin (ALB) (**i-j**). In (**h**), a marked increase in collagen is seen in ARPKD organoids. CK8 counterstaining indicates the marked increase in the size and number of ductal structures within the ARPKD organoids. In (**h-i**), the ducts in control organoids have a simple columnar CK19^+^ epithelium (indicated by arrows), while the ductal epithelium in ARPKD organoids is thickened (indicated by arrowheads) and abnormal. Scale bars are 50 μm.

The basis for the abnormal ductular morphology was investigated by immunofluorescence staining. In control organoids, zonula occludens protein 1 (ZO-1) and EZRIN were expressed in a characteristic manner on the apical side of the cholangiocytes surrounding the ductal lumen (**Fig. 2a).** This pattern indicates that ductal epithelial cells formed tight junctions that were properly oriented with respect to the ductal plane ^43^, which explains why the control organoids had a normal tubular architecture. In contrast, ZO-1 expression was decreased in ARPKD organoids, and was present in a non-oriented manner within the ductal structures. Also, the characteristic expression pattern of a cell polarity-determining protein (Vang-Like 1, VANGL1) in CK19^+^ cholangiocytes, which is oriented around ductal structures in control organoids, was similarly altered in ARPKD organoids (**Fig. 2b**). These results indicate that the orientation and polarity of bile duct cholangiocytes was maintained in control organoids; but these features were disrupted in the ARPKD organoids. Also, while primary cilium are formed in both control and ARPKD organoids, they are ~2-fold more abundant in ARPKD (vs control) organoids (**Fig. 2c**). Thus, the ARPKD mutation does not interfere with the formation of primary cilium, but it could impact their function.

**Figure 2.**
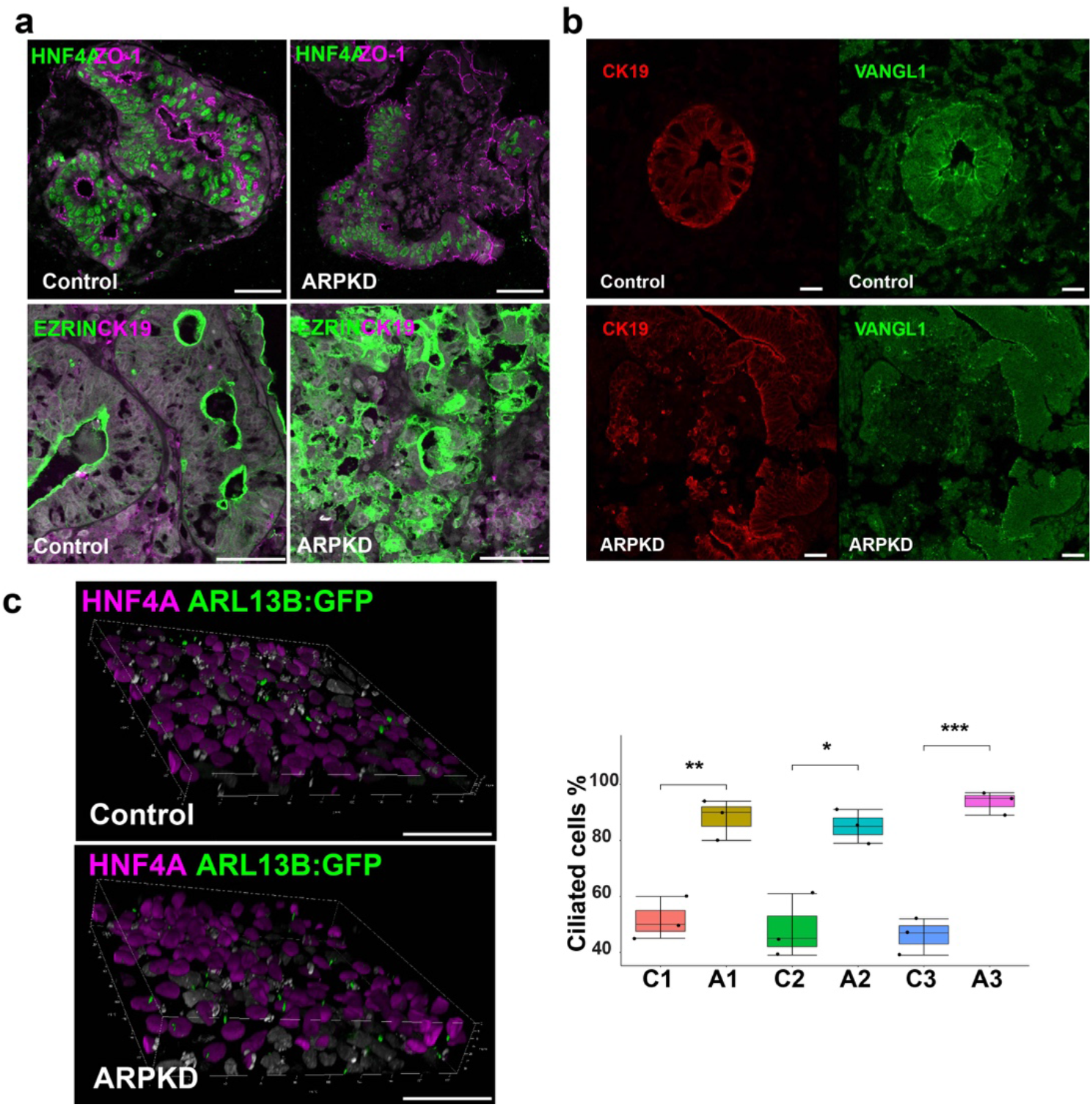
Loss of polarity and increased number of primary cilia in ARPKD organoids. **a**, HOs were immunostained with antibodies to ZO-1 and HNF4A, EZRIN and CK19; While control organoids have a localized pattern of ZO-1 and EZRIN expression that surrounds the apical side of duct lumen; ZO-1 and EZRIN expression in ARPKD organoids is diffuse and is not oriented around the ducts. **b**, HOs were immunostained with antibodies to VANGL1 and CK19. VANGL1 is highly expressed in CK19^+^ cells in control HOs; but its expression is decreased, and it is expressed more diffusely in the cytoplasm of cells in ARPKD organoids. **c**, *Left:* 3D stacked confocal images of ARL13B:GFP fusion protein expression in control and ARPKD organoids. *Right:* A graph showing the percentage of cells with primary cilium in control (C1, C2, C3) and ARPKD organoids (A1, A2, A3). These box plots show the average, 25-75% range and SE of measurements made on >100 stacked images for each set of three organoids. *, p value <0.05; **, p, value <0.01; and ***, p value <0.001. While primary cilium are formed in both control and ARPKD organoids, they are more abundant in ARPKD than in control organoids. All scale bars are 50 μm.

### Quantitative assessment of fibrosis in ARPKD HOs and ARPKD liver tissue

Liver tissue obtained from ARPKD subjects had enlarged bile ducts, and a markedly increased amount of ECM (including collagen fibers) was deposited throughout the liver tissue (**Fig. 3a**). Hence, ARPKD organoid morphology mirrors the major pathologic features observed in ARPKD liver disease, which includes abnormal bile ducts and increased ECM deposition that is characteristic of CHF. As described in the supplemental note (Fig. S1d), second harmonic generation (**SHG**) microscopy has been used to analyze liver fibrosis ^44, 45^. Therefore, 3D SHG images were obtained to quantitatively evaluate collagen fiber abundance and structure within unlabeled control and ARPKD hepatic organoids after 21 days of *in vitro* culture. Whereas the crosslinked collagen in control organoids was assembled into thin (diameter<1.5 μm) fibers that surrounded the cells in a few, isolated regions; ARPKD organoids had a confluent network of thick collagen fibers throughout the entire organoid (**Fig. 3b**). A quantitative analysis confirmed the marked increase in cross-linked collagen fibers in ARPKD organoids (average volume fraction 17.0%+ 6.8%, n=43 organoids) relative to control organoids (average volume fraction 3.2%+2.9%, n=40 organoids); and the increase was noted irrespective of whether ARPKD organoids prepared from the 3 donors were evaluated individually (p<0.05) or as a mixture (p<0.0001) relative to isogenic controls (Fig. 3b). Moreover, the abundance of thicker collagen fibers (diameter ≥ 6.0 μm) in ARPKD organoids was significantly greater than in isogenic control organoids (individual p<0.05, mixture p<0.001).

**Figure 3.**
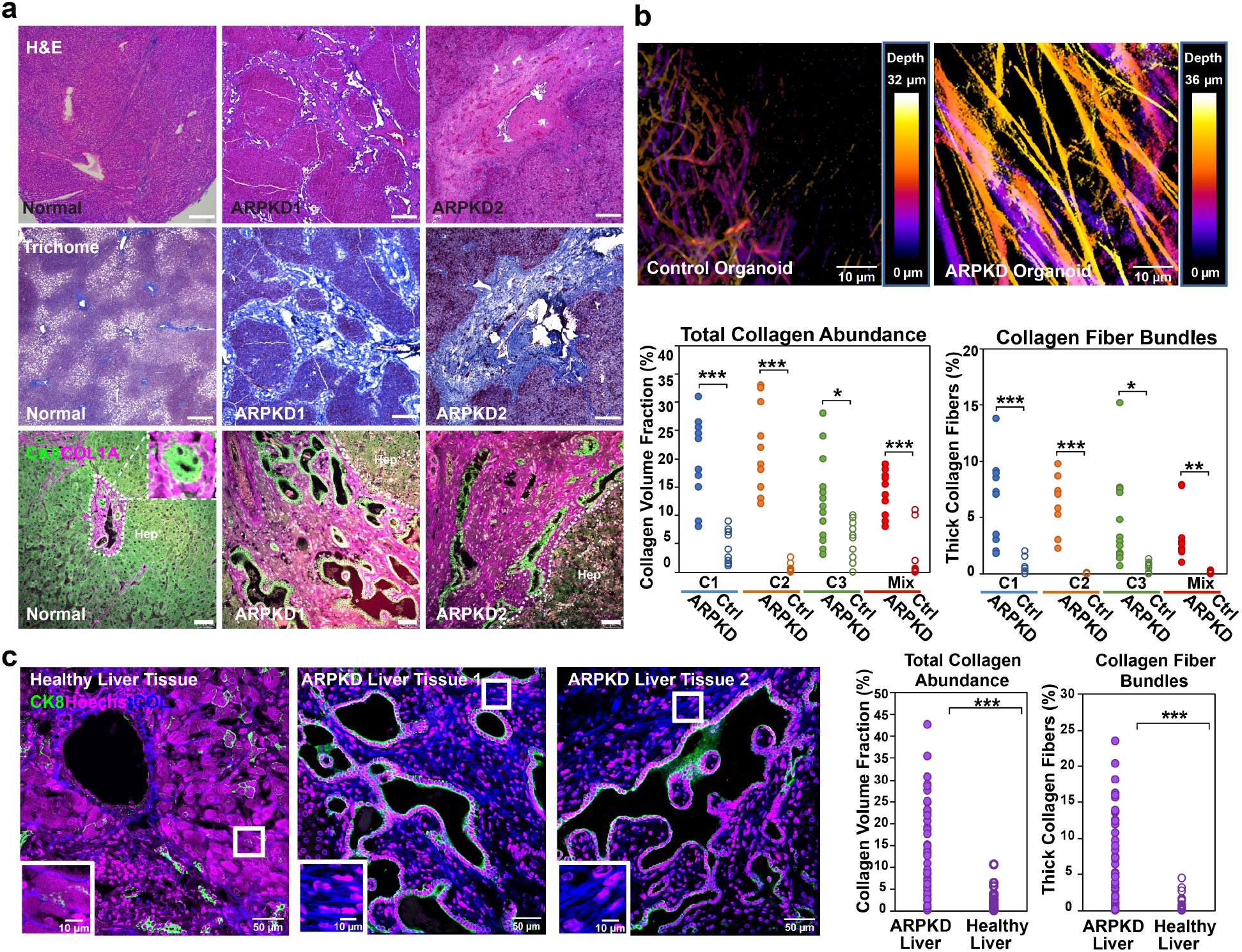
ARPKD liver tissue has enlarged bile ducts and a marked increase in collagen. Liver sections were prepared from a normal subject and from two subjects with ARPKD liver disease. **a**, In the H&E (**upper**) and trichrome-stained liver sections (middle), the marked increase in ECM (blue regions) in ARPKD liver tissue is readily apparent. Scale bars: 500 μm. Bottom panels: Higher power (scale bars, 50 μm) immunofluorescent images of liver sections stained with anti-CK8 and anti-collagen (COL1A) antibodies. CK8 is a marker for both hepatocytes and cholangiocytes. An area with hepatocytes is shown in the normal liver tissue. In normal liver, some collagen is deposited around the portal triads; and the inset shows an enlarged view of a portal triad. In ARPKD liver samples, there is a marked increase in the amount of collagen, which is diffusely distributed throughout the stroma; and the abnormal bile ducts lined by CK8^+^ cells are readily apparent. **b,** SHG microscopic analysis of the collagen fibers in human hepatic organoids. (**upper**) Depth color-coded projections of collagen fibers within day 21 control and ARPKD organoids. Control organoids (left) have a few, isolated regions with relatively thin collagen fibers. ARPKD organoids (right) have a confluent network of thick collagen fibers that extend throughout the entire organoid, both laterally as well as in depth. (**lower**) Quantitative comparison of SHG images from ARPKD and control organoids (N ≥ 10 for C1, C2, C3; and for a mixed culture of all three) shows statistically significant increases in total collagen abundance (left), and in the fraction of thick collagen fiber bundles (right, diameter > 6.0 μm). Unpaired *t*-test results (ARPKD vs control): *p < 0.05, **p < 0.001, ***p < 0.0001. **c**, SHG analysis of liver tissue obtained from control and two ARPKD subjects. In these images, the collagen fibers are blue; DAPI-stained nuclei are magenta colored; and CK19^+^ cholangiocytes are green. The amount of fibrous collagen is significantly increased in the ARPKD liver samples (13.3 + 10.6%, n=47) compared with normal liver tissue (1.8 + 2.3%, n=38). Unpaired *t*-test results (ARPKD vs healthy liver): ***p < 0.0001. Also, the collagen fibers in the ARPKD liver tissue formed thick bundles (right, diameter > 6.0 μm, unpaired *t*-test ARPKD vs health liver ***p < 0.0001), which are much larger than the ~1 micron-sized, isolated fibers present in the normal liver tissue (see insets).

To determine if the ARPKD organoids mirrored ARPKD liver pathology, liver tissue samples obtained from normal and ARPKD patients were analyzed by SHG microscopy. Similar to what was observed in the organoids, the collagen fiber volume in the ARPKD liver tissue (13.3% + 10.6%) was much greater than in control liver tissue (1.7% + 1.2%; p =0.0001 **Fig. 3c**). Additionally, the collagen fibers in the ARPKD liver tissue formed thicker, aligned bundles that were much larger than the ~1 micron-sized, isolated fibers present in normal liver tissue. A significantly higher volume fraction of thick collagen bundles (diameter ≥ 6.0 μm) was present in ARPKD tissue (p < 0.0001). Of note, we did not observe fluid filled cysts in ARPKD organoids, nor in the ARPKD liver tissue examined here. Usually, ARPKD patients do not develop polycystic liver disease, which is a finding that is characteristic of Autosomal Dominant Polycystic Kidney Disease (ADPKD). In contrast, ARPKD liver disease is characterized by congenital fibrosis and ductal abnormalities, as was observed in the ARPKD organoids. Overall, the SHG analyses indicated that the ARPKD mutation caused a marked increase in collagen fiber abundance and dimensions; and that the pathologic changes observed in ARPKD organoids mirrored those occurring in ARPKD liver tissue.

### Pathogenesis of ARPKD liver disease

A multiplex analysis of scRNA-Seq data generated from ARPKD and isogenic control HOs prepared from 3 unrelated individuals (C1-C3) was performed ^46^ (**Figs. S4a**). In total, the transcriptomes of 7,461 cells in control organoids (average 61,000 reads and 2,884 genes per cell), and of 11,960 cells in the corresponding ARPKD organoids (average 36,000 reads and 2,079 genes per cell) were analyzed (**Figs. S4b**). A cell clustering and dimensional reduction analysis identified 15 clusters in control and ARPKD organoids, which was visualized using the t-Distributed Stochastic Neighbor Embedding (**t-SNE**) method (**Figs. 4a-b, S4c**). While each of the different cell types was tightly clustered, there were significant differences in the cellular composition of control and ARPKD organoids. The transcriptome of cell clusters in the hepatic organoids were compared with cells in control and cirrhotic human livers ^47^, and the initial analysis indicated that various types of mesenchymal cells (clusters 0,1, 3, 4, 6, 7), mesothelial cells (cluster 2), hepatocyte precursor cells (cluster 5), early endothelia (clusters 8, 9), cholangiocytes (cluster 10), and endothelia (cluster 14) were present in the organoids (**Fig. 4a**, **Table S1).** Cluster identities were confirmed by examination of the levels of expression of mRNAs encoding canonical markers **(Fig. S4d**). To identify cells that could contribute to ARPKD liver pathology, the frequency of cells in each cluster within control and ARPKD organoids was calculated (**Fig. 4a**). Multiple clusters (5, 8, and 12-14) had a similar percentage of cells in control and ARPKD organoids, which indicates that changes the percentage of cells observed for the other clusters were not caused by a preparation artifact. However, there was a dramatic shift in the mesenchymal populations present in ARPKD and control organoids. The percentage of cells within clusters 0, 1 and 7 were 9-, 7- and 7.8-fold increased, respectively, in ARPKD organoids relative to control organoids; while the percentage of clusters 3 and 4 were 10- and 5-fold increased, respectively, in control organoids; and cluster 6 was present in control organoids (14.5%) but was virtually absent (<0.1%) in ARPKD organoids. Thus, there was an increase in the number and type of mesenchymal cells present in ARPKD organoids. Consistent with the immunostaining results (Fig. 2a-b), the expression levels for multiple mRNAs associated with the primary cilium or with planar cell polarity were decreased in ARPKD HOs relative to isogenic control HOs (**Fig. S4e).**

**Figure 4.**
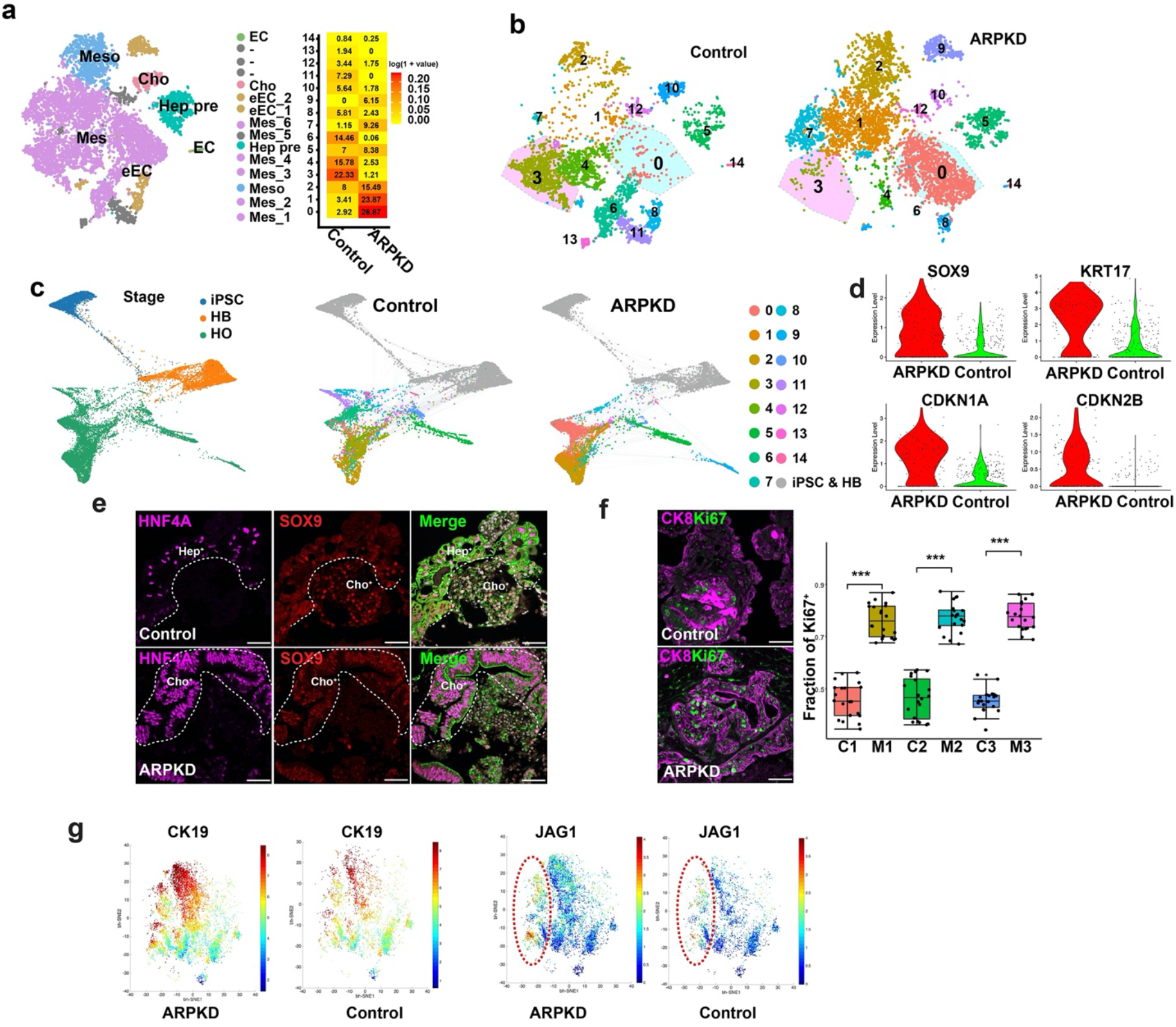
scRNA-Seq analysis of hepatic organoids. **a**, A combined t-SNE plot of the ARPKD and control organoid scRNA-Seq data shows the cell types identified within the 15 different clusters that were identified by k-nearest neighbor analysis within the Seurat suite, which include: various types of mesenchymal cells (Mes; clusters 0, 1, 3, 4, 6, 7), mesothelia (Meso; cluster 2), hepatocyte precursors (Hep pre; cluster 5), early endothelia (eEC, clusters 8, 9), cholangiocytes (Cho; cluster 10), and endothelia (EC; cluster 14). The heatmap (**on right**) compares the percentages of cells within each cluster that are present in control and ARPKD organoids. The colors represent the log transformed percentage of cells within each cluster. The cell types within the 15 clusters are listed in **Table S1,** and were annotated as described in **Table S2.** The percentage of cells within clusters 0 and 1 are 9- and 7-fold increased, respectively, in ARPKD (vs. control) organoids; while clusters 3 and 4 are 10- and 5-fold increased, respectively, in control (vs ARPKD) organoids. **b**, t-SNE plots separately show the 15 clusters identified in control and ARPKD organoids. **c**, The developmental trajectories of ARPKD and isogenic control cells are similar at the iPSC (day 0) and hepatoblast (HB, day 9) stages, but differ significantly at the organoid (day 21) stage. The **left** graph shows the cells in ARPKD and isogenic control cultures at the iPSC, HB and hepatic organoid stages. The ARPKD and isogenic control cells are separately graphed (**middle, right**). Each of the 15 different cell clusters in the organoids is indicated by a color; while the iPSC and HB cells are gray. The ARPKD and isogenic control cells are very similar at the iPSC and HB stages, but significantly diverge at the organoid stage. As examples, clusters 0 and 1 are far more abundant in ARPKD than in control organoids; while clusters 3 and 4 are far more abundant in control organoids; and cluster 9 is present in ARPKD, but not in control organoids. **d**, Violin plots show the increased level of expression of mRNAs for liver progenitor cell (SOX9, 2.1-fold; KRT17, 5.3-fold) and cell proliferation (CDKN1A 3.2-fold; CDKN2B, 1.9-fold) markers in ARPKD relative to control cholangiocytes (cluster 10). **e-f,** Control and ARPKD HOs were immunostained with antibodies to SOX9 and HNF4A or with anti-CK8 and anti-Ki67 antibodies. The dashed lines separate areas with hepatocytes (Hep^+^) or cholangiocytes (Cho^+^) from other regions. The boxplot shows the fraction of Ki67^+^ cholangiocytes within 3 pairs of ARPKD and isogenic control HOs. **g,** CK19 and JAG1 expression is increased in ARPKD organoids. bhSNE maps of CyToF data generated using ARPKD and control HOs. The dotted circle shows the JAG1^+^ cell population.

To identify the developmental stage when the ARPKD mutation alters the differentiation trajectory of cells within the organoid cultures, we analyzed scRNA-Seq data generated from ARPKD and isogenic control iPSCs, hepatoblasts, and organoid cultures prepared from the 3 unrelated individuals. In total, the transcriptomes of 10,000 ARPKD and 10,000 isogenic control cells were analyzed. The developmental trajectories of ARPKD and isogenic control cells are quite similar at the iPSC (day 0) and hepatoblast (day 9) stages, but significantly differed at the organoid stage (**Fig. 4c, S5a-e**). We also analyzed previously obtained scRNA-Seq data ^39^ to determine when the primary cilium genes, which are mutated in ARPKD or ADPKD (*PKD1*), are expressed during organoid development. *PKHD1* mRNA is expressed in iPSCs, but its expression level was significantly decreased at the hepatoblast stage, and then increased at the organoid stage. *PKD1* mRNA is expressed at the hepatoblast and HO stages **(Fig. S5f).** RT-PCR analyses indicated that equivalent levels of the mRNAs encoding four mesenchymal cell markers (*COL1A1, PDGFRB, Vim, ACA2*) were expressed in day 9 ARPKD and control hepatoblast cultures (**Fig. S5g**). Taken together, these results indicate that the ARPKD mutation-induced effect on mesenchymal populations probably occurs during the period when hepatoblasts differentiate into the cells that are present the mature liver organoid.

### ARPKD mutation effects on cholangiocytes

To investigate the pathogenesis of ARPKD ductal abnormalities, we compared the transcriptomes of the cholangiocytes (cluster 10) in ARPKD and control organoids. The level of expression of 439 mRNAs were altered in ARPKD cholangiocytes relative to control cholangiocytes (Fold Change > 1.3, minimum fraction of cells >0.1). The 10 most differentially expressed genes identified by this comparison are shown in **Table S2**. Reactome Pathway Database (https://reactome.org/) analysis identified 62 pathways that were significantly enriched among the 439 genes whose expression level was increased in ARPKD cholangiocytes (**Fig. S6a**). Of note, the most highly enriched pathway was extracellular matrix organization (**Fig. S6b,** fold enrichment 14.1, lowest p-value 8.4 x 10^−9^); and *MMP2, COL1A2, TIMP2* and *MMP9* mRNAs were 2.6, 5.2, 1.5 and 1.7-fold-increased in ARPKD cholangiocytes (**Fig. S6c**). ARPKD cholangiocytes had an increased level of expression of multiple mRNAs that regulate the cell cycle and mitosis, collagens and proteins involved in collagen fiber assembly, and of genes expressed in liver progenitor cells (**Fig. 4d, Fig. S6d**). Immunostaining shows that ARPKD HOs have an abnormal ductular morphology, which is characterized by a thickened and disoriented layer of cells around the ductal lumen. Of note, while the cholangiocytes in ARPKD organoids are HNF4A and SOX9 double positive cells; cholangiocytes in control organoids are either HNF4A or SOX9 positive, but these mRNAs are not co-expressed in the same cells (**Fig. 4e**). Ki67 staining shows that ARPKD HOs have many more proliferating cholangiocytes than control organoids (**Fig. 4f**). CyTOF analysis also showed that JAG1^+^ cells, which are CK19^+^ cholangiocytes, were 1.5-fold more abundant in ARPKD than in control HOs (**Fig. 4g**). Thus, ARPKD cholangiocytes are more proliferative; they produce enzymes and other proteins that promote a high level of collagen bundle generation; and they are less mature than control cholangiocytes. Of particular interest because of its known role in the pathogenesis of fibrosis, the transcriptome of ARPKD cholangiocytes were enriched for mRNAs within TGFβ-associated signaling pathways (TGFβ signaling in EMT, signaling by TGFβ Receptor Complex). Also, ARPKD cholangiocytes had a 1.3-fold increased level of TGFβ1 mRNA expression (p=3.1×10^−8^) and a 4.8-fold increase in the level of a TGFβ inducible mRNA (*TGFBI*) (**Fig. S6e).** ARPKD cholangiocytes also express reduced levels of mRNAs encoding planar cell polarity (*DVL2, FZD6*) and ductal epithelium/tight junction (*CLDN1, CLDH1, TJP1, EPCAM*) proteins than control organoids (**Fig. S6f**). Thus, transcriptome analysis indicates that ARPKD cholangiocytes are less mature, have a reduced level of expression of mRNAs regulating cell polarity and epithelial cell function; but have increased TGFβ pathway activation, and are more actively involved in collagen fiber generation than control cholangiocytes.

### Myofibroblast cell expansion in ARPKD

Of the 15 identified cell clusters, cluster 0 was of particular interest because: (i) it had the largest increase in cell number in ARPKD (27%) versus control (2.9%) organoids; (ii) and pathway enrichment analysis indicated that they expressed mRNAs associated with protein digestion, the JAK-STAT pathway, and ECM-receptor interactions (**Fig. 5a-c**); Most importantly, (iii) of the six mesenchymal cell clusters present in hepatic organoids, the cluster 0 transcriptome was most similar to that of myofibroblast-like cells found in cirrhotic human liver tissue ^47^ (**Fig S7a).** Moreover, of the 15 cells clusters, cluster 0 cells had the highest level of *PDGFRB* mRNA, which is myofibroblast marker **(Fig. S7b**). Immunostaining and CyTOF results indicated that ARPKD organoid cells had a marked increase in PDGFRβ, SMA, PDGFRa and VIM protein expression; and the percentage of PDGFRβ^+^ cells in ARPKD organoids was 4.5-fold increased (versus control organoids) (**Figs. 5d-f, S7c**). While a small number of SMA^+^ cells were occasionally present in control organoids; ARPKD organoids had an increased number of SMA^+^ cells, which were present in clusters located near ductal structures (**Fig. 5f**). These organoid features were reflected in the markedly increased level of PDGFRβ and SMA protein expression observed in ARPKD liver tissue (**Fig. 5g**). Thus, by multiple criteria, the cluster 0 transcriptome resembles that of myofibroblasts; and their number was markedly increased in ARPKD organoids.

**Figure 5.**
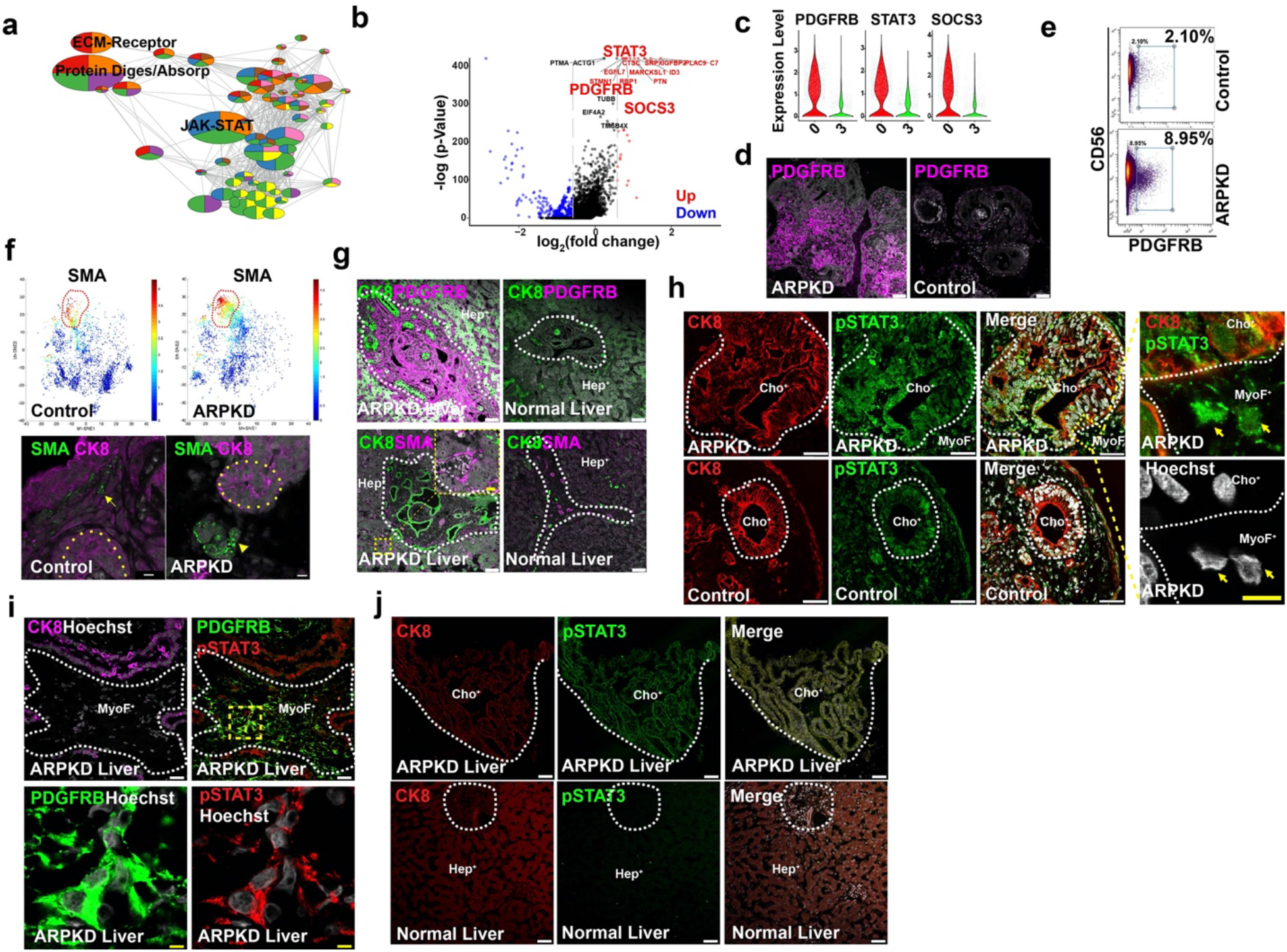
ARPKD hepatic organoids have an expanded population of myofibroblast cells that resemble those in fibrotic human liver tissue. **a**, KEGG pathway enrichment analysis identifies gene networks whose expression is increased in cluster 0 cells. The gene networks involved in protein digestion and adsorption, the JAK-STAT signaling pathway, and extracellular matrix (ECM)-receptor interactions were most highly enriched within cluster 0 cells. **b**, A volcano plot showing the differentially expressed genes (fold change > 1.5, adjust p-value < 0.05,) when the transcriptomes of cluster 0 and cluster 3 are compared. The 20 genes whose mRNAs exhibit the highest level of differential expression are highlighted, and these include STAT3, PDGFRβ and SOCS3. The gray dashed line indicates a 1.5-fold change, which is the cutoff for a differentially expressed gene. A red (or blue) color indicates that a gene is upregulated (or down regulated) in cluster 0 (vs. cluster 3); a black color indicates that the expression level was not significantly different. **c**, Violin plots showing three genes (*PDGFRB, STAT3, SOCS3*) whose mRNAs were markedly increased in cluster 0 cells relative to cluster 3. **d**, Day 21 control and ARPKD organoids were immunostained with an anti-PDGFRβ antibody. PDGFRB is highly expressed in ARPKD, but not in control, hepatic organoids (HO). Scale bar, 50 μm. **e**, A scatter plot of CyToF data performed with anti-PDGRFβ and anti-CD56 antibodies shows that ARPKD organoids have a markedly increased amount of PDGFRB^+^ cells (9.0%) relative to control organoids (2.1%). **f**, bhSNE maps (**upper**) generated from CyToF data indicate that ARPKD HOs have an increased amount of SMA^+^ cells. (**Lower**) Representative images of HOs immunostained with anti-smooth muscle actin (SMA) and anti-CK8 antibodies. The yellow dotted lines indicate ductal structures. While only a few SMA^+^ cells were present at a limited number of sites within control organoids (indicated by an arrow); large clusters of SMA^+^ cells were present within ARPKD organoids (indicated by an arrowhead), and were located near ductal structures. Scale bar, 10 μm. **g**, Normal and ARPKD liver tissue was immunostained with antibodies to PDGFRβ, CK8 and SMA. PDGFRβ expression was much higher in ARPKD than in control liver tissue. Also, SMA^+^ cells were only located within perivascular regions of normal liver tissue; but they infiltrate the parenchyma of ARPKD liver tissue (as shown in the inset). The white dashed line indicated epithelium and mesenchymal part. Scale bar, 50 μm. **h**, ARPKD and Control HOs were immunostained with anti-CK8 and anti-phospo-STAT3 (Tyr705) antibodies. Phospho-STAT3 was extensively expressed in bile duct and mesenchymal cells in ARPKD HOs. The images shown in the right panels are enlargements of the boxed yellow area indicated in the ARPKD organoid; the arrows indicate mesenchymal cells with Phospho-STAT3 within the nucleus. Scale bar, 5 μm. **i**-**j**, ARPKD and normal liver tissues were immunostained with antibodies to CK8, Phospho-Stat3 and CK8, and were counterstained with Hoechst. (**i**) Low (top) and high (bottom) power views of ARPKD liver tissue immunostained with antibodies to CK8, PDGFRB and Phospho-STAT3. An area with myofibroblasts is enclosed within the dotted line. A high-power view of the yellow box area shows phospho-STAT3 is co-localized within PDGFRB^+^ myofibroblasts. Scale bar, 5 μm. (**j**) Phospho-STAT3 is extensively expressed in an enlarged bile duct within ARPKD liver tissue, but its level of expression is much weaker expression in normal bile ducts.

While different mesenchymal cell populations were present in control and ARPKD organoids, the myofibroblast-like cluster 0 cells that were most abundant in ARPKD organoids (27% of total) were distinctly different from cluster 3 cells that were the most abundant in control organoids (22% of total). Of particular interest, relative to cluster 3, cluster 0 cells have increased levels of mRNAs for multiple JAK-STAT3 signaling pathway components and for its effector molecules such as PIM-1 ^48, 49^ and *PDGFRβ*, and for multiple RNAs that regulate stem cell pluripotency and fibroblast activation (**Fig. S8a-b**). For example, cluster 0 cells had an increased level of *leukemia inhibitory factor receptor (LIFR*) mRNA, which is of interest because LIF induces an invasive and activated state in fibroblasts in a STAT3-dependent manner ^50^. *LIF* mRNA was expressed at a low level within multiple different cell clusters, which was similar to its pattern in human liver tissue; and clusters 0 and 3 expressed equivalent *LIF* mRNA levels (**Fig. S8c**). Similarly, *PDGFA* and *PDGFB* mRNAs were expressed by multiple cell types within the organoids (**Fig. S8c**). In contrast, cluster 3 cells had increased levels of mRNAs associated with cell cycle arrest and/or cellular senescence [i.e. *TP53* ^51^, GADD45b ^52^] (**Fig. S8d).** Thus, the ARPKD mutation promotes the production of myofibroblast-like cells that have the characteristics of activated and proliferative cells, and their transcriptome is quite distinct from the mesenchymal cells in control organoids. Also, *STAT3* mRNA expression increased during HO differentiation (**Fig. S8e**); and the active phosphorylated form of STAT3 was present in ARPKD HO myofibroblasts and in ARPKD liver tissue. Interestingly, Phospho-STAT3 was also found in cholangiocytes in ARPKD organoids and in areas with abnormal ducts in ARPKD liver tissue (**Fig. 5h-j**).

### Similarities with commonly occurring forms of human liver fibrosis

We wanted to investigate whether the ARPKD organoid fibrosis mechanistically resembled that in the commonly occurring forms of human liver fibrosis. Transcriptome analysis identified 254 genes whose expression was increased in the myofibroblast-like cells (cluster 0) in the ARPKD organoid, which were used to form a myofibroblast-specific expression signature (**Table S3**). Gene Set Enrichment Analysis (**GSEA**) ^53^ was used to assess whether this myofibroblast expression signature was present in other types of fibrotic liver tissue. GSEA has been used to identify genes/pathways associated with treatment response or disease prognosis ^54–56^, and to identify stem cell signatures in human cancer tissues ^57, 58^. GSEA calculates a normalized expression score (**NES**), which indicates whether myofibroblast signature genes are enriched in fibrotic liver tissue. GSEA analysis was performed using expression data obtained from 10 normal and 10 hepatitis C virus infection-induced cirrhotic liver tissues (GSE6764, ^59^). The myofibroblast expression signature was very strongly associated with cirrhotic liver (NSE 2.56, false discovery rate (FDR) 0), but not normal liver (NES −2.55, FDR 0) (**Fig. S9a-b**). We next investigated whether the myofibroblast signature was associated with non-alcoholic steatohepatitis (**NASH**), which is now the most common cause of chronic liver disease ^60, 61^. Although NASH is triggered by an abnormal triglyceride accumulation; fibrosis develops and progresses as NASH liver disease advances. Myofibroblast activation is key to its pathogenesis ^8, 62, 63^, and the extent of liver fibrosis is the major determinant of NASH outcome ^64, 65^. Therefore, a gene expression dataset (GSE83452) containing 98 normal and 126 NASH liver tissues was analyzed. The myofibroblast expression signature was strongly associated with NASH liver (NSE 1.65, FDR 0), but not with normal liver tissue (NES −1.64, FDR 0). Of importance, in the absence of liver fibrosis, the myofibroblast expression signature was not induced by obesity (NES 0.98, FDR 0.55) or hepatocellular carcinoma (NES 0.24, FDR 0.4) (**Fig. S9a**). We next directly compared the transcriptome of myofibroblasts in ARPKD organoids with myofibroblasts present in cirrhotic human livers ^47^. The myofibroblast gene signature of cirrhotic liver tissue was strongly correlated with ARPKD but not with control organoids (**Fig. S9b**). As controls, GSEA results indicated that a B cell signature was not significantly present in either ARPKD or control organoids; and that the hepatocyte and cholangiocyte expression signatures were evenly distributed between control and ARPKD organoids (**Fig. S9c**). Thus, two different types of GSEA analyses indicate that the myofibroblasts in ARPKD organoids resemble those that cause the commonly occurring (acquired) forms of human liver fibrosis.

### Inhibitor effect on fibrosis

Our transcriptomic and other analyses indicated that the PDGFRB/STAT3 pathway was highly activated in ARPKD organoids. To determine if this pathway was essential to the pathogenesis of ARPKD fibrosis, we examined the effect that three clinically used PDGFR inhibitors (Crenolanib ^66, 67^, Sunitinib ^68, 69^ and Imatinib ^70, 71^) had on the extent of ARPKD organoid fibrosis. Three different methods were used to assess collagen abundance in ARPKD organoids: quantitative collagen immunostaining in whole organoids, and measurement of hydroxyproline and collagen mRNA levels. Immunostaining and analysis of whole-mount organoids indicated that treatment with 10 uM concentrations of PDGFR (Imatinib, Crenolanib) or NOTCH (DAPT) inhibitors significantly decreased collagen formation in ARPKD organoids (p<0.001 vs untreated organoids) (**Fig. 6a**). In fact, the fibrosis scores in the drug-treated ARPKD organoids were close to that of control hepatic organoids (p-value = 0.068 control vs drug-treated ARPKD organoids) (**Fig. 6b**). To determine if the inhibitors affected mRNA transcription, we examined their effect on *COL1A* mRNA levels in ARPKD organoids. *COL1A1* mRNA levels in ARPKD organoids were decreased ~17-fold after treatment with the three PDGFR inhibitors (p < 0.001) (**Fig. 6c**). We also measured 4-hydroxyproline (4-OH Pro) levels, which increase in fibrotic tissue ^72^, in the vehicle and drug-treated ARPKD organoids. Consistent with their effect on *COL1A* mRNA and on collagen fiber formation, all three PDGFRB inhibitors at 10 uM (p<0.005) and 0.05 uM (p<0.005) concentrations significantly decreased 4-OH Pro levels in ARPKD organoids (**Fig. 6d**). The anti-fibrotic effect of the PDGFR pathway inhibitors confirms that the PDGFRB pathway contributes to the development of fibrosis in ARPKD organoids.

**Figure 6.**
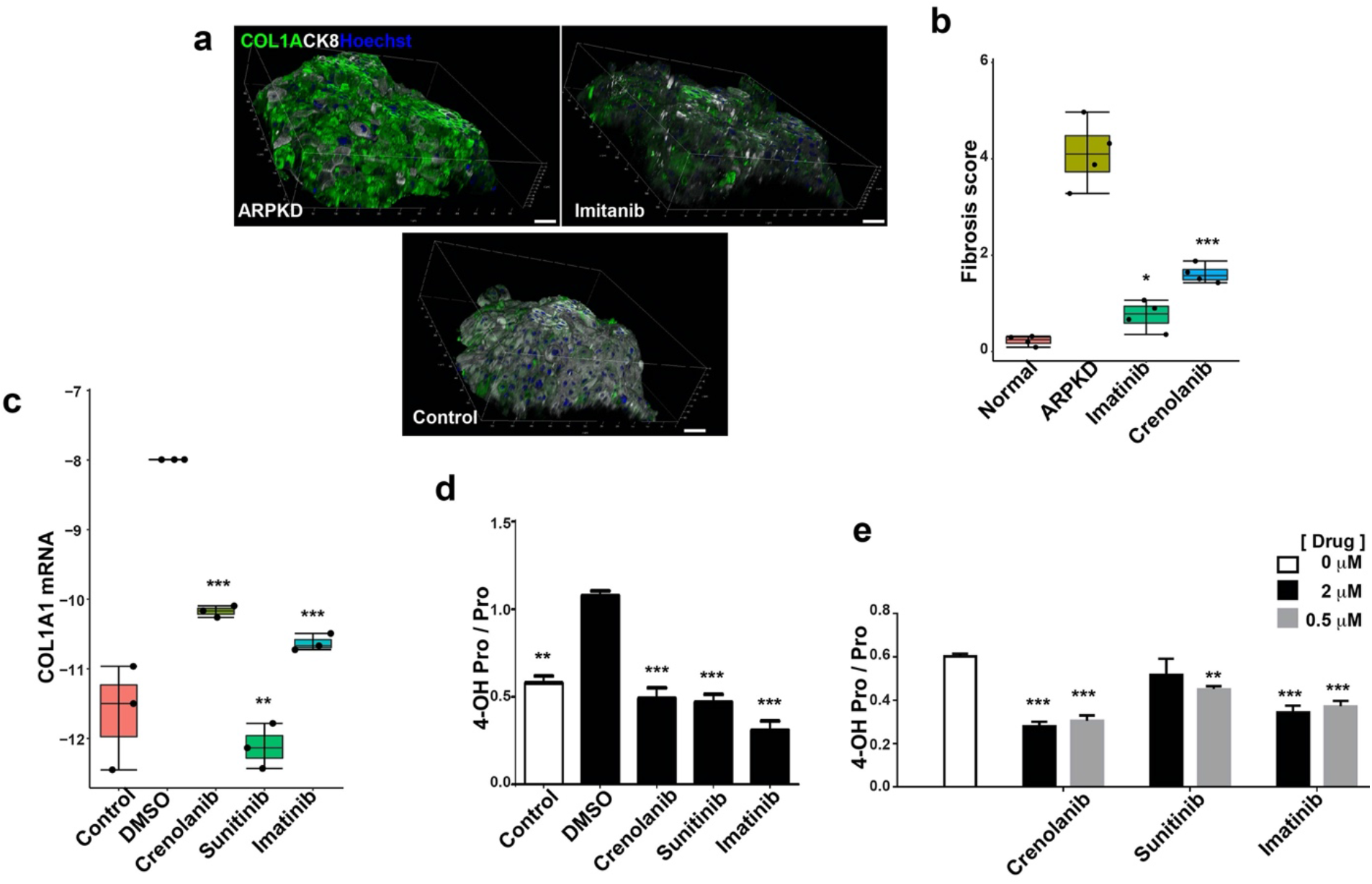
Inhibitors of the PDGFR and NOTCH signaling pathways reduce fibrosis in ARPKD organoids. Control or ARPKD organoid cultures were treated with vehicle or with PDGFR inhibitor from days 12 through 21, and the extent of fibrosis developing in the cultures was measured by three different methods. (**a**) Whole-mount immunostaining was performed on control HOs, and on ARPKD organoids that were treated with vehicle or 10 uM of the indicated drug. Representative images of organoids that were stained with antibodies to COL1A and CK8, and counterstained with Hoechst 33342 to show the nuclei. Scale bar, 50 μm. (**b**) The fibrosis score was determined by measuring the ratio of the volume of COL1A and CK8 in 100 stacked images obtained from 4 wholemount organoids per group. Each box plot shows the average and the 25% and 75% values for each group. The fibrosis score is significantly greater in vehicle-treated ARPKD organoids than in control organoids. Treatment with the PDGFR inhibitors (Crenolanib, Imatinib) decreased the fibrosis score (p<0.01) in ARPKD organoids to levels that were not significantly different from the control organoids (p>0.05). (**c**) *COL1A1* mRNA levels in ARPKD organoids are significantly higher than in control organoids; and *COL1A1* mRNA levels in ARPKD organoids were decreased 17-fold after treatment with the PDGFR (Crenolanib, Sunitinib and Imatinib) inhibitors (p < 0.001). Each box plot shows the average, and the 25% and 75% for RT-PCR measurements made on 3 organoids per treatment group; and the values were normalized relative to simultaneously measured *GAPDH* mRNA levels. (**d,e**) Four-hydroxyproline levels (4-OH Pro) in ARPKD organoids are decreased by treatment with 10 μM (d), 2 μM or 0.5 μM (e) concentrations of PDGFRB inhibitors. Each bar shows the average of the 4-OH Pro levels + SEM (shown as the ratio of the 4-OH Pro to proline levels) measured in 3 control or ARPKD organoids treated with either vehicle or the indicated concentration of each drug. In d: the 4-OH Pro levels in control organoids (control) are significantly below that of the ARPKD organoids (**, p<0.0005); and the 4-OH Pro levels in drug-treated ARPKD organoids are significantly below those of the vehicle-treated (0 uM drug) ARPKD organoids (***, p<0.0001). In e, 4-OH Pro levels were measured in vehicle (0 uM) or drug-treated ARPKD organoids. The 4-OH Pro levels in the drug-treated ARPKD organoids that are significantly below the vehicle-treated ARPKD organoids are indicated (***, p<0.002; **, p<0.01)

## Discussion

ARPKD liver pathology, which includes bile duct abnormalities and fibrosis, develops in hepatic organoids with an engineered mutation in *PKHD1* (which encodes a mutated FPC) that is the most common cause of ARPKD. The widely discordant phenotypes appearing within some families has led to suggestions that genetic modifiers could affect ARPKD disease expression ^15, 73^. However, ARPKD pathology developed in organoids independently of donor genetic background, and comparison with isogeneic controls confirmed that the pathology is induced by the ARPKD mutation. Also, since bile fluid does not flow in the organoid cultures, mutation-induced abnormalities in the mechanosensory function of primary cilium do not appear to contribute to the pathogenesis of ARPKD liver disease. A detailed analysis of ARPKD and isogenic control HOs indicates that four mutation-induced alterations are essential to the pathogenesis of ARPKD liver disease: (i) ARPKD cholangiocytes are less mature; they produce an increased amount of active TGF-β; their transcriptional program is strongly altered by TGF-β-associated signaling; and they appear to be actively involved in collagen thick fiber generation; (ii) the ARPKD mutation promotes the expansion of collagen-producing myofibroblasts, which are distinguished by their having an activated STAT3 signaling pathway and a markedly increased level of PDGFRβ expression; (iii) the amount and dimensions of the collagen fibers are increased; and (iv) the orientation and polarity of ductal cholangiocytes is disrupted.

Activation of the TGF-β-associated signaling pathway in the cholangiocytes in ARPKD organoids is consistent with prior *in vitro* and *in vivo* observations. ARPKD cholangiocytes in rodent models have an increased level of expression of TGF-β1 and of its receptor ^74, 75^. In cultured cells, the ARPKD mutation disrupts the interaction between FPC and the NEDD4 family member ubiquitin E3 ligase complex; and this enhances TGF-β signaling by impairing the degradation of the TGF-β receptor ^76^. Moreover, the FPC mutation in cholangiocytes have an increase in *MMP-2* and *MMP-9* mRNAs that could increase the conversion of latent TGF-β into its active form ^77, 78^, which could further amplify the effect of the ARPKD mutation on TGF-β signaling. STAT3 pathway activation in ARPKD myofibroblasts is consistent with evidence suggesting that this pathway plays a role in ADPKD, since it can be augmented by a mutation in a membrane protein (polycystin-1, PC1) that forms a complex with FPC ^79^. The increased level of expression of STAT3 pathway-associated mRNAs (*Myc, PDGFRβ, PIM-1*) in ARPKD myofibroblasts is also of interest. The marked increase in PDGFRβ protein expression is consistent with the well-known role that the PDGFR/STAT pathway has in promoting hepatic fibrogenesis ^80^, and PDGFRβ cross-linking leads to STAT3 phosphorylation and activation ^81^. Myc, which is induced by PDGF in a STAT3-dependent manner ^82^, promotes the proliferation of hepatic stellate cells and their conversion into myofibroblasts ^83^. ARPKD myofibroblasts also had increased *SOCS3* mRNA levels, which, under normal conditions downregulates the STAT3 signaling pathway ^84, 85,86,87,88^. ARPKD myofibroblasts also have an increased level of *LIF receptor* mRNA expression, and they (along with cholangiocytes and possibly other cell types) produce LIF (**Fig. S8**). The LIF receptor forms a cell membrane-localized complex with gp130 ^89^, whose expression is down-regulated by SOCS3 ^88^. Although ARPKD fibroblasts have increased *SOCS3* mRNA levels, this “brake” on the system, which would normally reduce both gp130 (as an inducer of the STAT3-stimulating pathway) and downstream elements thereof ^84–89^ could be overwhelmed by the combined activation of STAT3 via PDGFRβ and LIF receptor signals. Whereas TGF-β1 alone was not able to induce cultured rodent ARPKD cholangiocytes to differentiate into mesenchymal cells ^74^, it acts in concert with LIF to induce (in a STAT3-dependent manner) fibroblasts to develop into cells with activated and invasive properties ^50^. This information along with our organoid data can be assembled into a potential model for the pathogenesis of ARPKD liver disease (**Fig. S10**). In brief, the ARPKD mutation in *PKHD1* generates cholangiocytes that produce an increased amount of TGF-β1, as well as the mesenchymal cell-derived enzymes and proteins involved in thick collagen fiber generation. The TGF-β1 produced by ARPKD cholangiocytes acts in concert with LIF and downstream phospho-STAT3 to jointly stimulate mesenchymal cells to become activated myofibroblasts. As depicted in this model, TGFβ and STAT3 signaling pathways are known to interact ^90, 91^. Ligand binding by TGFβ receptors activates the signal transducing SMAD proteins ^92^. SMAD binding sites are often located near the STAT3 binding sites (downstream of LIF), and these genomic regions (known as ‘enhanceosomes’) play an important role in defining cellular identity ^93^. This could generate a self-sustaining circuit that acts in conjunction with STAT3 pathway activation-associated effects – which include the increased level of expression of *Myc, Fos, Jun, PDGFRβ* and other mRNAs in myofibroblasts - to generate and maintain the fibrotic state. The ability of PDGFRβ inhibitors to significantly reduce the extent of fibrosis in ARPKD organoids confirms that the PDGFRβ-STAT3 pathway plays an important role in the pathogenesis of liver fibrosis. Additional investigations are required to confirm some of the other aspects of this model.

This is the first liver organoid system that models the key feature of human liver fibrosis, which is the formation of an excess of assembled collagen fibers. We establish that the fibrosis in ARPKDs results from an increased abundance of thick collagen fibers and we demonstrate that its features mirror that of fibrotic human liver tissue. Moreover, the multi-lineage cells within our model develop in a coordinated fashion to produce an organoid with liver features. Although the fibrosis studied here is genetically induced, myofibroblasts are the key mediators of all types of liver fibrosis, including the commonly occurring forms that are caused by chronic alcohol exposure or viral diseases ^7, 94–96^. Some evidence indicates that common mechanisms could underlie fibrotic diseases of diverse etiologies occurring in different tissues ^97^. Irrespective of whether liver fibrosis develops in response to an acquired or congenital cause, aside from treatments aimed at the underlying cause, no available treatments can prevent or reverse its progression. A major limitation on therapeutic development has been the model systems used for studying fibrosis. Prior to the emergence of organoid methodology, *in vitro* cellular models of fibrosis were limited by several factors. They often used plated hepatocytes; which lack the three dimensional architecture of liver, and they do not have the different cell types involved in fibrogenesis ^98^. Most models require some type of injury-induced activation ^98, 99^, which induces variability in the response. Commonly used rodent models of liver fibrosis also require an injury inducing agent (carbon tetrachloride, bile duct ligation ^100^, or dietary modulation), and they are time consuming and expensive to run. Since fibrosis is an intrinsic feature of the ARPKD organoid model, no exogenous disease-inducing agent is required, which eliminates the variability that is introduced by the injury-inducing agent. Moreover, any conclusions drawn from animal models are limited by concerns about interspecies differences and about their fidelity with the processes mediating human liver fibrosis. In contrast, this hepatic organoid model is a human-based *in vitro* system that generates a 3-dimensional multi-lineage liver-like tissue with bile ducts, and it has all of the enzymes required to synthesize collagen and to form cross-linked collagen fibers. Furthermore, the myofibroblast gene expression signature present in ARPKD organoids resembles that found in the commonly occurring acquired forms of human fibrotic liver diseases. These attractive features indicate that use of this hepatic organoid system could increase our ability to produce new treatments for fibrotic liver disease.

## Supporting information

Sup Info ARPKD rev

## Acknowledgements

We thank Dr. Hyun Min Kang for his generous advice on the use of the demuxlet program for deconvoluting multiplexed scRNA-Seq data; and Dr. Bob Lewis for helpful advice and thoughtful review of this manuscript.

## Abbreviations

ARPKD: Autosomal Recessive Polycystic Kidney Disease
CARS: coherent anti-Stokes Raman scattering
CHF: congenital hepatic fibrosis
CyTOF: high-dimensional time of flight mass cytometry
DAPT: N-[N-(3,5-Difluorophenacetyl)-L-alanyl]-S-phenylglycine t-butyl ester
ECM: extracellular matrix
EMT: epithelial to mesenchyme transition
FPC: fibrocystin/polyductin
GSEA: Gene signature expression analysis
HB: hepatoblast
HSC: Hepatic stellate cell
HO: hepatic organoid
iPSC: induced pluripotent stem cell
*PKHD1*: *polycystic kidney and hepatic disease-1*
SAMe: scar associated mesenchymal cells
SHG: Second Harmonic Generation
SMA: smooth muscle actin
t-SNE: t-Distributed Stochastic Neighbor Embedding

## References

1. Yoon YJ, Friedman SL, Lee YA. Antifibrotic Therapies: Where Are We Now? Semin Liver Dis 2016;36:87–98.

2. Hernandez-Gea V, Friedman SL. Pathogenesis of liver fibrosis. Annu Rev Pathol 2011;6:425–56.

3. Gourdie RG, Dimmeler S, Kohl P. Novel therapeutic strategies targeting fibroblasts and fibrosis in heart disease. Nat Rev Drug Discov 2016;15:620–38.

4. Rock N, Kanavaki I, McLin VA. Congenital Hepatic Fibrosis, Caroli’s Disease, and Other Fibrocystic Liver Diseases. In: Guandalini S, Dhawan A, Branski D, eds. Textbook of Pediatric Gastroenterology, Hepatology and Nutrition. New York: Springer International Publishing, 2016:647–662.

5. Pinzani M, Luong TV. Pathogenesis of biliary fibrosis. Biochim Biophys Acta Mol Basis Dis 2018;1864:1279–1283.

6. Pinzani M. Pathophysiology of Liver Fibrosis. Dig Dis 2015;33:492–7.

7. Zhang CY, Yuan WG, He P, et al. Liver fibrosis and hepatic stellate cells: Etiology, pathological hallmarks and therapeutic targets. World J Gastroenterol 2016;22:10512–10522.

8. Tsuchida T, Friedman SL. Mechanisms of hepatic stellate cell activation. Nat Rev Gastroenterol Hepatol 2017;14:397–411.

9. Yin C, Evason KJ, Asahina K, et al. Hepatic stellate cells in liver development, regeneration, and cancer. J Clin Invest 2013;123:1902–10.

10. Zhang DY, Goossens N, Guo J, et al. A hepatic stellate cell gene expression signature associated with outcomes in hepatitis C cirrhosis and hepatocellular carcinoma after curative resection. Gut 2016;65:1754–64.

11. Mederacke I, Hsu CC, Troeger JS, et al. Fate tracing reveals hepatic stellate cells as dominant contributors to liver fibrosis independent of its aetiology. Nat Commun 2013;4:2823.

12. Aycock RS, Seyer JM. Collagens of normal and cirrhotic human liver. Connect Tissue Res 1989;23:19–31.

13. Olaso E, Ikeda K, Eng FJ, et al. DDR2 receptor promotes MMP-2-mediated proliferation and invasion by hepatic stellate cells. J Clin Invest 2001;108:1369–78.

14. Schuppan D, Ruehl M, Somasundaram R, et al. Matrix as a modulator of hepatic fibrogenesis. Semin Liver Dis 2001;21:351–72.

15. Hartung EA, Guay-Woodford LM. Autosomal recessive polycystic kidney disease: a hepatorenal fibrocystic disorder with pleiotropic effects. Pediatrics 2014;134:e833–45.

16. Buscher R, Buscher AK, Weber S, et al. Clinical manifestations of autosomal recessive polycystic kidney disease (ARPKD): kidney-related and non-kidney-related phenotypes. Pediatr Nephrol 2014;29:1915–25.

17. Wilson PD. Polycystic kidney disease. N Engl J Med 2004;350:151–64.

18. Jiang L, Fang P, Weemhoff JL, et al. Evidence for a “Pathogenic Triumvirate” in Congenital Hepatic Fibrosis in Autosomal Recessive Polycystic Kidney Disease. Biomed Res Int 2016;2016:4918798.

19. Hildebrandt F, Benzing T, Katsanis N. Ciliopathies. N Engl J Med 2011;364:1533–43.

20. Ward CJ, Hogan MC, Rossetti S, et al. The gene mutated in autosomal recessive polycystic kidney disease encodes a large, receptor-like protein. Nat Genet 2002;30:259–69.

21. Onuchic LF, Furu L, Nagasawa Y, et al. PKHD1, the polycystic kidney and hepatic disease 1 gene, encodes a novel large protein containing multiple immunoglobulin-like plexin-transcription-factor domains and parallel beta-helix 1 repeats. Am J Hum Genet 2002;70:1305–17.

22. Bergmann C, Senderek J, Windelen E, et al. Clinical consequences of PKHD1 mutations in 164 patients with autosomal-recessive polycystic kidney disease (ARPKD). Kidney Int 2005;67:829–48.

23. Bergmann C, Senderek J, Kupper F, et al. PKHD1 mutations in autosomal recessive polycystic kidney disease (ARPKD). Hum Mutat 2004;23:453–63.

24. Melchionda S, Palladino T, Castellana S, et al. Expanding the mutation spectrum in 130 probands with ARPKD: identification of 62 novel PKHD1 mutations by sanger sequencing and MLPA analysis. J Hum Genet 2016.

25. Bergmann C, Kupper F, Dornia C, et al. Algorithm for efficient PKHD1 mutation screening in autosomal recessive polycystic kidney disease (ARPKD). Hum Mutat 2005;25:225–31.

26. Bergmann C. Genetics of Autosomal Recessive Polycystic Kidney Disease and Its Differential Diagnoses. Front Pediatr 2017;5:221.

27. Harris PC, Rossetti S. Molecular genetics of autosomal recessive polycystic kidney disease. Mol Genet Metab 2004;81:75–85.

28. Kim I, Fu Y, Hui K, et al. Fibrocystin/polyductin modulates renal tubular formation by regulating polycystin-2 expression and function. J Am Soc Nephrol 2008;19:455–68.

29. Wang S, Zhang J, Nauli SM, et al. Fibrocystin/polyductin, found in the same protein complex with polycystin-2, regulates calcium responses in kidney epithelia. Mol Cell Biol 2007;27:3241–52.

30. Rohatgi R, Battini L, Kim P, et al. Mechanoregulation of intracellular Ca2+ in human autosomal recessive polycystic kidney disease cyst-lining renal epithelial cells. Am J Physiol Renal Physiol 2008;294:F890–9.

31. Banales JM, Masyuk TV, Gradilone SA, et al. The cAMP effectors Epac and protein kinase a (PKA) are involved in the hepatic cystogenesis of an animal model of autosomal recessive polycystic kidney disease (ARPKD). Hepatology 2009;49:160–74.

32. Spirli C, Locatelli L, Morell CM, et al. Protein kinase A-dependent pSer(675) -beta-catenin, a novel signaling defect in a mouse model of congenital hepatic fibrosis. Hepatology 2013;58:1713–23.

33. Guan Y, Xu D, Garfin PM, et al. Human Hepatic Organoids for the Analysis of Human Genetic Diseases. JCI Insight 2017;2:pii: 94954.

34. Villani AC, Satija R, Reynolds G, et al. Single-cell RNA-seq reveals new types of human blood dendritic cells, monocytes, and progenitors. Science 2017;356.

35. Zeisel A, Munoz-Manchado AB, Codeluppi S, et al. Brain structure. Cell types in the mouse cortex and hippocampus revealed by single-cell RNA-seq. Science 2015;347:1138–42.

36. Montoro DT, Haber AL, Biton M, et al. A revised airway epithelial hierarchy includes CFTR-expressing ionocytes. Nature 2018;560:319–324.

37. Plasschaert LW, Zilionis R, Choo-Wing R, et al. A single-cell atlas of the airway epithelium reveals the CFTR-rich pulmonary ionocyte. Nature 2018;560:377–381.

38. Halpern KB, Shenhav R, Matcovitch-Natan O, et al. Single-cell spatial reconstruction reveals global division of labour in the mammalian liver. Nature 2017;542:352–356.

39. Guan Y, Chen X, Wu M, et al. The phosphatidylethanolamine biosynthesis pathway provides a new target for cancer chemotherapy. J Hepatol 2019;72:746–760.

40. Furu L, Onuchic LF, Gharavi A, et al. Milder presentation of recessive polycystic kidney disease requires presence of amino acid substitution mutations. J Am Soc Nephrol 2003;14:2004–14.

41. Quint A, Sagi M, Carmi S, et al. An Ashkenazi founder mutation in the PKHD1 gene. Eur J Med Genet 2016;59:86–90.

42. Kilpinen H, Goncalves A, Leha A, et al. Common genetic variation drives molecular heterogeneity in human iPSCs. Nature 2017;546:370–375.

43. Huang RY, Guilford P, Thiery JP. Early events in cell adhesion and polarity during epithelial-mesenchymal transition. J Cell Sci 2012;125:4417–22.

44. Sun W, Chang S, Tai DC, et al. Nonlinear optical microscopy: use of second harmonic generation and two-photon microscopy for automated quantitative liver fibrosis studies. J Biomed Opt 2008;13:064010.

45. Gailhouste L, Le Grand Y, Odin C, et al. Fibrillar collagen scoring by second harmonic microscopy: a new tool in the assessment of liver fibrosis. J Hepatol 2010;52:398–406.

46. Kang HM, Subramaniam M, Targ S, et al. Multiplexed droplet single-cell RNA-sequencing using natural genetic variation. Nat Biotechnol 2018;36:89–94.

47. Ramachandran P, Dobie R, Wilson-Kanamori JR, et al. Resolving the fibrotic niche of human liver cirrhosis at single-cell level. Nature 2019;575:512–518.

48. Stout BA, Bates ME, Liu LY, et al. IL-5 and granulocyte-macrophage colony-stimulating factor activate STAT3 and STAT5 and promote Pim-1 and cyclin D3 protein expression in human eosinophils. J Immunol 2004;173:6409–17.

49. Shirogane T, Fukada T, Muller JM, et al. Synergistic roles for Pim-1 and c-Myc in STAT3-mediated cell cycle progression and antiapoptosis. Immunity 1999;11:709–19.

50. Albrengues J, Bourget I, Pons C, et al. LIF mediates proinvasive activation of stromal fibroblasts in cancer. Cell Rep 2014;7:1664–1678.

51. Aubrey BJ, Kelly GL, Janic A, et al. How does p53 induce apoptosis and how does this relate to p53-mediated tumour suppression? Cell Death Differ 2018;25:104–113.

52. Ou DL, Shen YC, Yu SL, et al. Induction of DNA damage-inducible gene GADD45beta contributes to sorafenib-induced apoptosis in hepatocellular carcinoma cells. Cancer Res 2010;70:9309–18.

53. Subramanian A, Tamayo P, Mootha VK, et al. Gene set enrichment analysis: a knowledge-based approach for interpreting genome-wide expression profiles. Proc Natl Acad Sci U S A 2005;102:15545–50.

54. Verstockt B, Verstockt S, Veny M, et al. Expression Levels of 4 Genes in Colon Tissue Might be Used to Predict Which Patients Will Enter Endoscopic Remission After Vedolizumab Therapy for Inflammatory Bowel Diseases. Clin Gastroenterol Hepatol 2019.

55. Wang Z, Wang Z, Niu X, et al. Identification of seven-gene signature for prediction of lung squamous cell carcinoma. Onco Targets Ther 2019;12:5979–5988.

56. Labrecque MP, Coleman IM, Brown LG, et al. Molecular profiling stratifies diverse phenotypes of treatment-refractory metastatic castration-resistant prostate cancer. J Clin Invest 2019;130.

57. Merlos-Suarez A, Barriga FM, Jung P, et al. The intestinal stem cell signature identifies colorectal cancer stem cells and predicts disease relapse. Cell Stem Cell 2011;8:511–24.

58. Corominas-Faja B, Oliveras-Ferraros C, Cuyas E, et al. Stem cell-like ALDH(bright) cellular states in EGFR-mutant non-small cell lung cancer: a novel mechanism of acquired resistance to erlotinib targetable with the natural polyphenol silibinin. Cell Cycle 2013;12:3390–404.

59. Davis S, Meltzer PS. GEOquery: a bridge between the Gene Expression Omnibus (GEO) and BioConductor. Bioinformatics 2007;23:1846–7.

60. Younossi ZM, Koenig AB, Abdelatif D, et al. Global epidemiology of nonalcoholic fatty liver disease-Meta-analytic assessment of prevalence, incidence, and outcomes. Hepatology 2016;64:73–84.

61. Sayiner M, Koenig A, Henry L, et al. Epidemiology of Nonalcoholic Fatty Liver Disease and Nonalcoholic Steatohepatitis in the United States and the Rest of the World. Clin Liver Dis 2016;20:205–14.

62. Sircana A, Paschetta E, Saba F, et al. Recent Insight into the Role of Fibrosis in Nonalcoholic Steatohepatitis-Related Hepatocellular Carcinoma. Int J Mol Sci 2019;20.

63. Marcher AB, Bendixen SM, Terkelsen MK, et al. Transcriptional regulation of Hepatic Stellate Cell activation in NASH. Sci Rep 2019;9:2324.

64. Tanaka N, Kimura T, Fujimori N, et al. Current status, problems, and perspectives of nonalcoholic fatty liver disease research. World J Gastroenterol 2019;25:163–177.

65. Angulo P, Kleiner DE, Dam-Larsen S, et al. Liver Fibrosis, but No Other Histologic Features, Is Associated With Long-term Outcomes of Patients With Nonalcoholic Fatty Liver Disease. Gastroenterology 2015;149:389–97 e10.

66. Smith CC, Lasater EA, Lin KC, et al. Crenolanib is a selective type I pan-FLT3 inhibitor. Proc Natl Acad Sci U S A 2014;111:5319–24.

67. Wang J, Fu X, Zhang D, et al. Effects of crenolanib, a nonselective inhibitor of PDGFR, in a mouse model of transient middle cerebral artery occlusion. Neuroscience 2017;364:202–211.

68. Huang X, Wang W, Yuan H, et al. Sunitinib, a Small-Molecule Kinase Inhibitor, Attenuates Bleomycin-Induced Pulmonary Fibrosis in Mice. Tohoku J Exp Med 2016;239:251–61.

69. Moran M, Nickens D, Adcock K, et al. Sunitinib for Metastatic Renal Cell Carcinoma: A Systematic Review and Meta-Analysis of Real-World and Clinical Trials Data. Target Oncol 2019;14:405–416.

70. Puckett RL, Lorey F, Rinaldo P, et al. Maple syrup urine disease: further evidence that newborn screening may fail to identify variant forms. Mol Genet Metab 2010;100:136–42.

71. Breedveld P, Pluim D, Cipriani G, et al. The Effect of Bcrp1 (Abcg2) on the In vivo Pharmacokinetics and Brain Penetration of Imatinib Mesylate (Gleevec): Implications for the Use of Breast Cancer Resistance Protein and P-Glycoprotein Inhibitors to Enable the Brain Penetration of Imatinib in Patients. Cancer Research 2005;65:2577–2582.

72. Gjaltema RA, Bank RA. Molecular insights into prolyl and lysyl hydroxylation of fibrillar collagens in health and disease. Crit Rev Biochem Mol Biol 2017;52:74–95.

73. Wills ES, Cnossen WR, Veltman JA, et al. Chromosomal abnormalities in hepatic cysts point to novel polycystic liver disease genes. Eur J Hum Genet 2016;24:1707–1714.

74. Sato Y, Harada K, Ozaki S, et al. Cholangiocytes with mesenchymal features contribute to progressive hepatic fibrosis of the polycystic kidney rat. Am J Pathol 2007;171:1859–71.

75. Moser M, Matthiesen S, Kirfel J, et al. A mouse model for cystic biliary dysgenesis in autosomal recessive polycystic kidney disease (ARPKD). Hepatology 2005;41:1113–21.

76. Kaimori JY, Lin CC, Outeda P, et al. NEDD4-family E3 ligase dysfunction due to PKHD1/Pkhd1 defects suggests a mechanistic model for ARPKD pathobiology. Sci Rep 2017;7:7733.

77. Perng DW, Chang, K.T., Su KC, et al. Matrix metalloprotease-9 induces transforming growth factorbeta(1) production in airway epithelium via activation of epidermal growth factor receptors. Life Sci 1989;89:204–212.

78. Kobayashi T, Kim H, Liu X, et al. Matrix metalloproteinase-9 activates TGF-beta and stimulates fibroblast contraction of collagen gels. Am J Physiol Lung Cell Mol Physiol 2014;306:L1006–15.

79. Talbot JJ, Shillingford JM, Vasanth S, et al. Polycystin-1 regulates STAT activity by a dual mechanism. Proc Natl Acad Sci U S A 2011;108:7985–90.

80. Ying HZ, Chen Q, Zhang WY, et al. PDGF signaling pathway in hepatic fibrosis pathogenesis and therapeutics (Review). Mol Med Rep 2017;16:7879–7889.

81. Vignais ML, Gilman M. Distinct mechanisms of activation of Stat1 and Stat3 by platelet-derived growth factor receptor in a cell-free system. Mol Cell Biol 1999;19:3727–35.

82. Bowman T, Broome MA, Sinibaldi D, et al. Stat3-mediated Myc expression is required for Src transformation and PDGF-induced mitogenesis. Proc Natl Acad Sci U S A 2001;98:7319–24.

83. Nevzorova YA, Hu W, Cubero FJ, et al. Overexpression of c-myc in hepatocytes promotes activation of hepatic stellate cells and facilitates the onset of liver fibrosis. Biochim Biophys Acta 2013;1832:1765–75.

84. Nicola NA, Nicholson SE, Metcalf D, et al. Negative regulation of cytokine signaling by the SOCS proteins. Cold Spring Harb Symp Quant Biol 1999;64:397–404.

85. Croker BA, Krebs DL, Zhang JG, et al. SOCS3 negatively regulates IL-6 signaling in vivo. Nat Immunol 2003;4:540–5.

86. Liau NPD, Laktyushin A, Lucet IS, et al. The molecular basis of JAK/STAT inhibition by SOCS1. Nat Commun 2018;9:1558.

87. Rui L, Yuan M, Frantz D, et al. SOCS-1 and SOCS-3 block insulin signaling by ubiquitin-mediated degradation of IRS1 and IRS2. J Biol Chem 2002;277:42394–8.

88. Kershaw NJ, Laktyushin A, Nicola NA, et al. Reconstruction of an active SOCS3-based E3 ubiquitin ligase complex in vitro: identification of the active components and JAK2 and gp130 as substrates. Growth Factors 2014;32:1–10.

89. Timmermann A, Kuster A, Kurth I, et al. A functional role of the membrane-proximal extracellular domains of the signal transducer gp130 in heterodimerization with the leukemia inhibitory factor receptor. Eur J Biochem 2002;269:2716–26.

90. Luo K. Signaling Cross Talk between TGF-beta/Smad and Other Signaling Pathways. Cold Spring Harb Perspect Biol 2017;9.

91. Chakraborty D, Sumova B, Mallano T, et al. Activation of STAT3 integrates common profibrotic pathways to promote fibroblast activation and tissue fibrosis. Nat Commun 2017;8:1130.

92. Hata A, Chen YG. TGF-beta Signaling from Receptors to Smads. Cold Spring Harb Perspect Biol 2016;8.

93. Chen X, Xu H, Yuan P, et al. Integration of external signaling pathways with the core transcriptional network in embryonic stem cells. Cell 2008;133:1106–17.

94. Higashi T, Friedman SL, Hoshida Y. Hepatic stellate cells as key target in liver fibrosis. Adv Drug Deliv Rev 2017;121:27–42.

95. Campana L, Iredale JP. Regression of Liver Fibrosis. Semin Liver Dis 2017;37:1–10.

96. Puche JE, Saiman Y, Friedman SL. Hepatic stellate cells and liver fibrosis. Compr Physiol 2013;3:1473–92.

97. Divakaruni SS, Van Dyke AM, Chandra R, et al. Long-Term Potentiation Requires a Rapid Burst of Dendritic Mitochondrial Fission during Induction. Neuron 2018;100:860–875 e7.

98. van Grunsven LA. 3D in vitro models of liver fibrosis. Adv Drug Deliv Rev 2017;121:133–146.

99. Mazza G, Al-Akkad W, Rombouts K. Engineering in vitro models of hepatofibrogenesis. Adv Drug Deliv Rev 2017;121:147–157.

100. Mariotti V, Cadamuro M, Spirli C, et al. Animal models of cholestasis: An update on inflammatory cholangiopathies. Biochim Biophys Acta Mol Basis Dis 2018.

