## Supplementary material for "A Human Multi-Lineage Hepatic Organoid Model for Liver Fibrosis": Sup Info ARPKD rev

*Supplemental note: Second Harmonic Generation (**SHG**) microscopy*. The spatial distribution and morphology of collagen fibers within organoids were monitored by SHG microscopy [1], [2]. SHG has been used to analyze human liver diseases that include fatty liver [3] and liver fibrosis [4, 5]. With the use of a high-numerical aperture objective, early signs of fibrosis (i.e. when submicron-sized collagen fibers start to aggregate) can be detected. Of importance, the information provided by SHG microscopy cannot be deduced by fluorescence microscopy of immunostained collagen, since immunostaining also detects non-fibrous collagen. Picrosirus red staining (imaged using polarization microscopy) is not capable of resolving thin collagen fibers; it requires thin-sectioning of samples so that no information can be provided on the 3D distribution of fibers; and it has been shown to produce data that is significantly different from that provided by the SHG signal [6]. SHG analysis revealed that control HOs had a network of thin cross-linked collagen fibers (average diameter< 1.5 $\mu$m) surrounding the cells in isolated regions. Coherent anti-Stokes Raman scattering (CARS) microscopy also showed that HOs had micron-sized intracellular lipid droplets (**Fig. S1d**). These results indicate that, besides synthesizing pro-collagen, cells within HOs have the enzymatic machinery required for cross-linking collagen to form fibers, as well as a functional lipid storing mechanism.

*Supplemental Note: Characterization of hepatic organoid differentiation*. scRNA-Seq analysis of iPSC, hepatoblast, and hepatic organoid cultures indicated that the differentiating cells could be separated into 5 distinct clusters that are consistent with their differentiation stage (**Figs. S2a-b**). In addition to the cells with canonical hepatocyte and cholangiocyte markers, cells expressing endothelial cell (PECAM1, ICAM1 and KDR), fibroblast (PDGFRB, COL1A1, DES and ACTA2) and possibly even macrophage (CD68, CD80, LGALS3 and PPARD) markers were also present (**Fig. S2c-f**). To plot the progression of iPSC to hepatic lineage differentiation, the “Monocle” package [7] was also used to construct single sell trajectories. It drew a branched single-cell trajectory beginning with iPSC, with a transition at the hepatoblast stage, and it terminated at the PHH and other cell types present in HO (**Fig. S2g**). A CyTOF analysis was performed to characterize the expression of multiple protein antigens on the cells at different differentiation stages within organoid cultures (**Fig. S2h**). The resulting bhSNE map clustered the cells into distinct populations, which included Tra1-81$^+$ iPS cells, CK19$^+$ hepatoblast cells, and CD26$^+$ hepatocytes and cholangiocytes (**Fig. S2h**). The protein expression signature also revealed that there was a continuous path of cellular differentiation within the HO cultures (**Fig. S2i**). We previously demonstrated that hepatic organoids could be dissociated, and that that individual cells could reform organoids that are referred to as secondary organoids (HO2) [8]. To do this, the dissociated cells are cultured in growth media (GM) for 6 days, and then in differentiation media for 6 days to produce the HO2. In some supplemental figures, the scRNA-Seq and CyTOF data analyzes HO2 cells cultured in GM or DM. Trichrome and immunostaining results indicate that HOs have a more complex pattern of antigen expression and form more complex structures than we previously appreciated. The organoids have ductal structures that are surrounded by mesenchymal cells and a collagen-enriched ECM; and anti-CD31 immunostaining indicates that they appear to form rudimentary vessels that are located away from ductal areas (**Fig. S3a-c**).

*Supplemental note: developmental effect of ARPKD mutation.* To identify the developmental stage affected by the ARPKD mutation, we analyzed scRNA-Seq data generated from ARPKD and isogenic control iPSCs, hepatoblasts, and organoid cultures prepared from the 3 unrelated individuals. In total, the transcriptomes of 10,000 ARPKD and 10,000 isogenic control cells were analyzed. The developmental trajectories of ARPKD and isogenic control cells are quite similar at the iPSC (day 0) and hepatoblast (day 9) stages, but significantly differed at the organoid stage (**Fig. 4c, S5a-e**). We also analyzed previously obtained scRNA-Seq data [9] to determine when the primary cilium genes, which are mutated in ARPKD or ADPKD (*PKD1*), are expressed during organoid development. *PKHD1* mRNA is expressed in iPSCs, but was significantly decreased at the hepatoblast stage, and then increased at the organoid stage. *PKD1* mRNA is expressed at the hepatoblast and HO stages **(Fig. S5f).** RT-PCR analyses indicated that equivalent levels of the mRNAs encoding four mesenchymal cell markers (*COL1A1, PDGFRB, Vim, ACA2*) were expressed in ARPKD and control hepatoblast cultures (**Fig. S5g**). Taken together, these results indicate that the ARPKD mutation-induced effect on mesenchymal populations probably occurs when hepatoblasts differentiate into the cells that are present the mature liver organoid.

*Supplemental Note: similarities with commonly occurring forms of human liver fibrosis*. To investigate whether the ARPKD organoid fibrosis mechanistically resembled that in the commonly occurring forms of human liver fibrosis, 254 genes whose expression was increased in the cluster 0 cells in the ARPKD organoid were used to form a myofibroblast-specific expression signature (**Table S3**). Gene Set Enrichment Analysis (**GSEA**) [10] was used to assess whether this myofibroblast expression signature was present in other types of fibrotic liver tissue. GSEA has been used to identify genes/pathways associated with treatment response or disease prognosis [11-13], and to identify stem cell signatures in human cancer tissues [14, 15]. GSEA calculates a normalized expression score (**NES**), which indicates whether myofibroblast signature genes are enriched in fibrotic liver tissue. GSEA analysis was performed using expression data obtained from 10 normal and 10 hepatitis C virus infection-induced cirrhotic liver tissues (GSE6764, [16]). The myofibroblast expression signature was very strongly associated with cirrhotic liver (NSE 2.56, false discovery rate (FDR) 0), but not normal liver (NES -2.55, FDR 0) (**Fig. S9a-b**). We next investigated whether the myofibroblast signature was associated with non-alcoholic steatohepatitis (**NASH**), which is now the most common cause of chronic liver disease [17, 18]. Although NASH is triggered by an abnormal triglyceride accumulation; fibrosis develops and progresses as NASH liver disease advances. Myofibroblast activation is key to its pathogenesis [19-21], and the extent of liver fibrosis is the major determinant of NASH outcome [22, 23]. Therefore, a gene expression dataset (GSE83452) containing 98 normal and 126 NASH liver tissues was analyzed. The myofibroblast expression signature was strongly associated with NASH liver (NSE 1.65, FDR 0), but not with normal liver tissue (NES -1.64, FDR 0). Of importance, in the absence of liver fibrosis, the myofibroblast expression signature was not induced by obesity (NES 0.98, FDR 0.55) or hepatocellular carcinoma (NES 0.24, FDR 0.4) (**Fig. S9a**). The myofibroblast gene signature of myofibroblast present in human cirrhotic liver tissue [24] was strongly correlated with ARPKD but not with control organoids (**Figs. S9b, 9c**). Thus, two different types of GSEA analyses indicate that ARPKD organoid myofibroblasts resemble those that cause the commonly occurring forms of human liver fibrosis.

*Supplemental comment: ARPKD model*. Activation of the TGF-β-associated signaling pathway in the cholangiocytes in ARPKD organoids is consistent with prior observations in ARPKD rodent models [25, 26]. In cultured cells, the ARPKD mutation disrupts the interaction between FPC and the NEDD4 family member ubiquitin E3 ligase complex; and this enhances TGF-β signaling by impairing TGF-β receptor degradation [27]. Moreover, an increase in *MMP-2* and *MMP-9* mRNAs in ARPKD cholangiocytes could increase the conversion of latent TGF-β into its active form [28, 29], which could further amplify the effect of the ARPKD mutation on TGF-β signaling. STAT3 pathway activation in ARPKD myofibroblasts is consistent with evidence suggesting that this pathway plays a role in ADPKD, which is caused by a mutated membrane protein (polycystin-1, PC1) that forms a complex with FPC [30]. Increased expression STAT3 pathway-associated mRNA expression (*Myc, PDGFRβ, PIM-1*) in ARPKD myofibroblasts is also of interest. The marked increase in PDGFRβ protein expression is consistent with the well-known role of the PDGFR/STAT pathway in promoting hepatic fibrogenesis [31], and PDGFRβ cross-linking leads to STAT3 phosphorylation and activation [32]. Myc, which is induced by PDGF in a STAT3-dependent manner [33], promotes the proliferation of hepatic stellate cells and their conversion into myofibroblasts [34]. ARPKD myofibroblasts also had increased *SOCS3* mRNA levels, which, under normal conditions downregulates the STAT3 signaling pathway [35, 36,37,38,39]. ARPKD myofibroblasts also have an increased level of *LIF receptor* mRNA expression, and they (along with cholangiocytes and possibly other cell types) produce LIF (Fig. S8). The LIF receptor forms a cell membrane-localized complex with gp130 [40], whose expression is down-regulated by SOCS3 [39]. Although ARPKD fibroblasts have increased *SOCS3* mRNA levels, this "brake" on the system, which would normally reduce both gp130 (as an inducer of the STAT3-stimulating pathway) and downstream elements thereof [35-40] could be overwhelmed by the combined activation of STAT3 via PDGFRβ and LIF receptor signals. Whereas TGF-β1 alone was not able to induce cultured rodent ARPKD cholangiocytes to differentiate into mesenchymal cells [25], it acts in concert with LIF to induce (in a STAT3-dependent manner) fibroblasts to develop into cells with activated and invasive properties [41]. This information along with our organoid data can be assembled into a potential model for the pathogenesis of ARPKD liver disease (**Fig. S10**). In brief, the ARPKD mutation in *PKHD1* generates cholangiocytes that produce an increased amount of TGF-β1, as well as the mesenchymal cell-derived enzymes and proteins involved in thick collagen fiber generation. The TGF-β1 produced by ARPKD cholangiocytes acts in concert with LIF and downstream phospho-STAT3 to jointly stimulate mesenchymal cells to become activated myofibroblasts. As depicted in this model, TGFβ and STAT3 signaling pathways are known to interact [42 43]. Ligand binding by TGFβ receptors activates the signal transducing SMAD proteins [44]. SMAD binding sites are often located near the STAT3 binding sites (downstream of LIF), and these genomic regions (known as 'enhanceosomes') play an important role in defining cellular identity [45]. This could generate a self-sustaining circuit that acts in conjunction with STAT3 pathway activation-associated effects – which include the increased level of expression of *Myc, Fos, Jun, PDGFRβ* and other mRNAs in myofibroblasts - to generate and maintain the fibrotic state.

### Materials and Methods

***iPSC and HO generation.*** Human iPSC lines (C1, C2 and C3) were prepared as previously described [8]. The methods for differentiating iPSCs into hepatic organoids (HOs) via culture in a series of media containing different growth factors were exactly as previously described [8].

***CRISPR-mediated genome engineering.*** CRISPR-associated protein 9 (Cas9) genome editing was coupled with the piggyBac transposon system to introduce the ARPKD mutation (*PKDH1 T36M)* into three different control (C1, C2 and C3) iPSC lines using our previously described method [8]. Site-specific (PKHD1 NG_008753.1 T36) guide RNA (sgRNA) sequences were selected using the CRISPR Design Tool [46]. Oligonucleotides with these sequences were cloned into the PX330 vector (Addgene, Cambridge, MA). The modifications of the piggyBac plasmid, which was originally obtained from the Sanger Institute, are described in [8]. For homology-directed repair (HDR), the piggyBac transposon was used with sequential culture media (containing puromycin and then ganciclovir) to select for cells with desired genome alterations as described [8]. To construct the mutation targeting vectors, two ~1 kb genomic fragments-each situated on one side of the *PKDH1 T36M* mutation-were PCR amplified and then cloned into the piggyBac transposon vector using the In-Fusion HD Cloning Kit (Clontech, Mountain View, CA). Engineered iPSCs were cloned and underwent two rounds of drug selection as previously described [8]. To confirm that the targeted mutations were correctly introduced, genomic DNA was isolated from the selected lines; PCR amplicons from the targeted loci were analyzed using the T7E1 enzyme assay; and were then cloned into a CloneJET PCR (Thermo Fisher Scientific, Grand Island, NY) cloning vector according to the manufacturer's instructions. Plasmid DNA isolated from 10 bacterial clones was analyzed by Sanger sequencing (Genewiz, San Francisco Lab, CA) to determine if homozygous alterations at the targeted site were introduced. All of the ARPKD iPSC used in this study were shown by sequencing to be homozygous *PKDH1 36M.*

***SHG and CARS microscopy***. The SHG (collagen fibers) and CARS (intracellular lipid stores) images of HOs were collected on a multimodal nonlinear optical microscope setup, consisting of a modified inverted confocal microscope (TE-2000, Nikon).  Two excitation beams from a tunable dual-beam near-infrared pico-second pulsed laser system (PicoEmerald, Applied Physics & Electronics, Inc., APE America) were overlapped in time and space, and coupled into a mirror scanner (C1/C2, Nikon) and focused by an oil immersion objective (CFI Apochromat, TIRF, 100X, numerical aperture (**NA**) 1.45, Nikon) for pixelwise scanning of the sample in x, y and z depth. The wavelengths of the excitation beams (pulse length: 2 ps, repetition rate: 80 MHz) were set to 797 nm and 1031 nm. Powers after the objective was 30 mW and 15 mW, respectively, measured with a microscope slide power sensor head. The SHG signal generated by the 797 nm beam was detected in back-reflection mode, optically filtered from the excitation beams by a dichroic beam splitter (735 nm edge, FF735-Di02-25x36, Semrock) and two filters (a 400/12 nm BrightLine single-band bandpass filter, FF01-400/12-25, and a 750 nm BrightLine Multiphoton SWP Filter, FF01-750/SP-25, Semrock). The combination of wavelengths (797 nm and 1031 nm) probes the carbon-hydrogen vibration (2845 $cm^{-1}$) of acyl chains in lipids. The corresponding coherent anti-Stokes Raman (CARS) signal from intracellular lipid stores was collected in transmission mode, simultaneously with the SHG signal, using the microscope condenser (NA 0.52, W.D. 30 mm) followed by optical filtering using two single-band bandpass filters (643/20 nm BrightLine single-band bandpass filter, FF01-643/20-25, Semrock). The following papers provide: a review of CARS microscopy[47]; a detailed description of CARS microscopy of intracellular lipid stores[48]; and describe its use for measurement of lipid stores in liver tissue [49, 50]. The CARS and SHG signals were each detected by an analog photomultiplier tube (Hamamatsu). The spatial resolution was estimated to ~0.3 $\mu$m in the x-y plane, and ~1 $\mu$m in depth.

Control and ARPKD HOs prepared from all donors from all were collected on day 21 fixed by incubation with a 4% paraformaldehyde solution and embedded in a droplet of media. They were then placed between two microscope cover slips (Fisher Scientific, #1) that were separated by a spacer with a circular hole that formed a well. In total, 84 SHG/CARS full volume images (z-stacks) were collected; 43 of ARPKD organoids and 41 of control organoids, n>10 per category (individual donor C1, C2, or C3, and mixed, respectively). The stacks covered a field-of-view of 149.9x149.9 $\mu$m, spanned by 1504x1504 pixels or 2048x2048 pixels. It resulted in a pixel size of ~100 nm or 73 nm. Each volume image (z-stack) was formed by 15-71 two-dimensional images. The pixel integration time was 1.2-2.4 $\mu$s. It is possible that hypoxia can develop within the central region of cultured organoids; and hypoxia is known to induce fibrosis [51]. Therefore, to exclude possible artifacts, which could be caused by hypoxia-induced expression fibrosis, the peripheral (<50 µm) regions of organoids were characterized in the SHG analyses. Images of normal and ARPKD human liver tissue were collected on a modified inverted Eclipse Ti2-E microscope (Nikon), which enabled the SHG results to be combined with confocal fluorescence microscopy.

All image processing and analysis was performed using the Fiji implementation of ImageJ [52]. Images were preprocessed by mean filtering (block size 5 pixels in x and y) and edges were highlighted using the FeatureJ Derivative plug-in. To quantify the volume fraction of collagen fibers within the probed sample, SHG images were binarized using the ImageJ Hysteresis thresholding plug-in. Two thresholds were defined to separate the images into three classes. All pixels with SHG signals below the lower threshold were set to zero; and all pixels with values above the high threshold were set to one. Pixels with intermediate values were set to one only if connected to a collection of pixels with values above the high threshold. This assured the separation of the full extent of the collagen fibers, despite few-pixel sized noise. The volume of the thresholded pixels (i.e. the collagen fibers) was calculated for each stack and divided by the total sample volume, forming the collagen volume fraction (in %). To determine the diameters of the collagen fibers, the fibers in the SHG images were quantified based upon their local thickness. The ImageJ Local Thickness routine (see the Analyze menu, Fiji) [53] was used, based on the Saito-Toriwaki Euclidean Distance Transformation Algorithm. The volume fraction of thick collagen fibers (diameter≥ 6 μm) was estimated by dividing the number of pixels represented by local thickness values ≥6 μm by the total pixel number of the stack. To test whether the volume fractions obtained for ARPKD and control organoids were significantly different, the unpaired Student's t-test was applied.

***scRNA-Sequencing.*** Hepatic organoids were prepared from each of the control iPSC lines (C1, C2, and C3) and from the corresponding iPSC with the *PKDH1 M36* mutation. The HO cultures were placed in a 1 ml solution containing 1 mg/ml collagenase and 1 mg/mg dispase (Worthington-Biochem, Lakewood, NJ), and then incubated for 15 to 30 minutes at 37°C with shaking. The cells were further dispersed by gentle pipetting with a fire-polished glass pipette, and then filtered through a 40 μm cell strainer (PluriSelect, San Diego, CA). The single cell suspension was visually inspected under a microscope, cells were counted using a Scepter™ 2.0 Handheld Automated Cell Counter (EMD çMillipore, Burlington, MA), and then resuspended in PBS with 0.01% BSA. Single cells were captured in droplets with barcoded beads using the 10x Genomics Chromium system (Pleasanton, CA). Cellular suspensions ($1x10^6$/ml) were then loaded onto the 10× Chromium instrument (10× Genomics); and were then sequenced according to the manufacturer's instructions. Three ARPKD and 3 control organoid samples were separately sequenced on two lanes of an Illumina HiSeq2500 Rapid Mode instrument, yielding 889M total reads.

***scRNA-sequencing for trajectory construction.*** iPSC, HB and HO cells were collected on days 0, 9 and 30, respectively, from isogenic control and ARPKD organoid cultures (from 3 different donors), and were prepared from according to previously described methods [8]. One million cells at each differentiation stage and group (ARPKD or isogenic control) were labeled using a total of 6 cell capture antibodies according to the manufacturer's protocol (BioLegend TotalSeq™). Then, the pooled cells were captured using two 10x Single Cell 3' Reagent Kit v3 runs; and sequenced on two lanes of an Illumina HiSeq2500 instrument, which yielded 660 M total reads that covered 10,000 cells in total.

***scRNA-Seq data analysis.*** Each sequencing lane generated data from 3 control or 3 ARPKD organoids. The datasets for each organoid was deconvoluted using the '*demuxlet*' method [54]. To do this, we analyzed previously obtained scRNA-Seq datasets [9], and identified 3.4M SNP alleles that could distinguish C1, C2 and C3 donors. These were used to generate a reference vCard file (VCF) that was analyzed by the demuxlet program. The donor for each cell within a lane was identified using the demuxlet program, which was run with the following parameters: the minor allele frequency was set to >1% to identify multiplets present in each dataset; we set '--alpha 0 --alpha 0.5', which assumes that the expected proportion of a 50% genetic mixture from two individuals; and we set '--group-list' to a list of 10X barcodes. By this method, the data in each lane was deconvoluted to determine the donor identity for each transcriptome.

After the identity of each cell was determined, an expression matrix was generated from 10x 'Cellranger', and the data was imported into 'Seurat' [55] for subsequent analysis. First, cells with unique gene counts <200 or where the percentage of mitochondrial mRNAs was >20% were removed. Then, a total of 4046 variable genes were identified using the default settings. Unwanted sources of variation were removed by regression analysis, which was performed to mitigate the effect of signals caused by mRNAs with unique molecular identifiers and mitochondria expression. Finally, a non-linear dimensional reduction (tSNE) program in 'Seurat' [55] was used to identify 15 unique clusters for all of the 12,580 cells analyzed. Fifteen principal components were used to construct the shared nearest neighbor (SNN) graph, and the parameter regulating the resolution of the ‘Find Clusters’ program, was set to 0.6.

PathfindR (*github: egeulgen/pathfindR*) was used to perform the pathway enrichment analysis through identification of active subnetworks. Differentially expressed genes (cluster biomarkers) identified from the Seurat analysis were used as the input genes. For cluster 0, 254 marker mRNAs were input for the analysis. In brief, a data frame consisting of the Gene Symbol, log-fold-change and adjusted-p values were used to perform the active subnetwork search. Pathway enrichment analyses was then performed using the genes identified within each of the identified active subnetworks using either the KEGG [56] or Reactome [57] database.

The scRNA-Seq data obtained from isogenic control and ARPKD mutated cells from all three donors at different stages of differentiation (iPSC, HB and organoid) were analyzed to evaluate their developmental trajectories. To do this, the cellular expression matrices were co-embedded using of the standard integrated workflow of the Seurat Program [58]. Thirty canonical components were used for the subsequent clustering analysis; and their differentiation trajectory was assessed using the Partition-based Graph Abstraction (PAGA) algorithm [59].

***Gene signature expression analyses (GSEA)***. The 455 differentially expressed mRNAs (455 genes) in ‘cluster 0’ were identified using the ‘*FindMarkers*’ function in ‘*Seurat*’ [55] with the parameter ‘min.pct’ set at 0.25 (which indicates the minimum fraction of cells within a group that expressed a given gene group of cells); and the default Wilcoxon rank sum test was used to perform this analysis. Of the 455 genes identified by this analysis, the expression level of 254 mRNAs were increased, while 201 mRNAs were decreased in ‘cluster 0.’ The myofibroblast expression dataset consisted of the 254 genes whose mRNAs were increased in ‘cluster 0.’ Three publicly available liver gene expression datasets (cirrhosis, GSE6764; NASH, GSE83452; Obesity, GSE126848), were obtained from the Gene Expression Omnibus using the ‘*GEOquery*’ as described [16]. Cirrhotic liver tissue was obtained from 10 subjects undergoing liver resection for hepatocellular carcinoma at one of 3 US or European hospitals; and their liver tissues were classified as cirrhotic by the examining pathologist. The hepatocellular carcinoma samples (HCC) samples examined in this study were obtained from eight subjects whose liver was resected for HCC, but these specimens did not have fibrosis or cirrhosis according to the examining pathologist. The 10 normal liver tissues (used for comparison) were obtained from 10 subjects undergoing liver resection for other reasons at the same hospitals, and their liver tissue was classified as normal by the examining pathologist [60]. For the NASH analysis, 231 liver biopsies were obtained from 87 subjects with varying degrees of abnormalities that were evaluated at the University of Antwerp. Based upon histological findings, which included the fibrosis stage and NAS score: 129 biopsies were classified as NASH, 98 as not having NASH, and 4 as indeterminate [61]. The 129 $NASH^+$ biopsies were compared with the 98 $NASH^-$ biopsies for this analysis. For the obesity comparison, liver tissue was obtained from 12 (otherwise healthy) obese individuals (BMI 30-40 $kg/m^2$) and from 14 healthy controls (BMI 18-25 $kg/m^2$), all without evidence of liver disease, who underwent a liver biopsy at the Center for Diabetes Research at the University of Copenhagen [62].

The myofibroblast signature GSEA was performed on the gene expression datasets generated from these samples according to previously described methods [10]; and 1000 permutations were used for significance assessment for each analysis. The Enrichment score (ES) reflects the degree to which the HSC gene set is overrepresented at the top or bottom of a ranked list of genes; the Normalized enrichment score (NES) was used to compare analysis results across gene sets; and the false discovery rate (FDR) was used to estimate the probability that a gene set with a given NES represents a false positive finding.

The gene signatures for the different cell types present in control and fibrotic human livers were obtained from [24]. Differentially expressed genes for pre-defined cell types were computed using the FindAllMarkers function within the Seurat Package [58]; and only those genes with a FDR < 0.05 and min.pct > 0.25 were assigned as cell type-specific signature genes. Then, the gene signatures for different types of liver cells was analyzed using the scRNA-Seq data generated from hepatic organoids.

***Transcriptome correlation.*** We assessed the correlation between the transcriptomes of mesenchymal cell clusters (0, 1, 3, 4, 6, and 7) in hepatic organoids with that of the four types of mesenchymal cells identified in control and cirrhotic human liver tissue in [24]: Mesenchyme 1, vascular smooth muscle cells; Mesenchyme 2, hepatic stellate cells; Mesenchyme 3, myofibroblasts (which were also referred to as scar associated mesenchymal cells (SAMe); and Mesenchyme 4, Mesothelia. To do this, the mean expression level for each gene in the cells within each cluster was determined, and these variable features were ranked based upon the number of clusters that they appeared in. The top 3000 features that were present in the organoid and liver tissue datasets were used for the correlation analysis.

***Patient samples and human ethics approval.*** Only de-identified ARPKD liver tissue was used in this study, which were obtained from Children's Hospital of Philadelphia (CHOP). Liver tissues were collected from 2 subjects with ARPKD liver disease that were treated at CHOP: ARPKD1: A05-56, 3 month fetus; and ARPKD2: CA09-04, age 34 weeks. The normal liver tissue used in this study was isolated from liver lobes that were resected due to liver cancer, and these samples were obtained with the approval of the Stanford University Medical Center IRB (IRB approval #42968). Regions with normal appearing tissue, which were spatially distant from the cancerous areas, were collected. Liver tissues were flash frozen on dry ice in Tissue-Tek® O.C.T™ (Sakura Finetek U.S.A., Torrance, CA). Tissue blocks were sectioned using a Leica CM3050 S Cryostat into 4 um sections, and serially obtained sections were stained.

***Histology and immunostaining.*** Fresh organoids were harvested, after settling by gravity, and were embedded in low melting point agarose (IBI Scientific, Dubuque, Iowa). Organoid blocks were then processed by sectioning of the paraffin-embedded tissue into 10-micron sections. The human liver tissues in 4 micron sections were fixed in 4% paraformaldehyde for 10 minutes, followed by permeabilization with 1% Triton X-

100 (Sigma-Aldrich, St. Louis, MO), and blocking was performed with a solution containing 10% chicken serum (Jackson ImmunoResearch, Bar Harbor, MA) for 30 min. Then, sections were incubated with the primary antibodies listed in **Table S4**. When needed, secondary antibody-staining was performed using an Alexa Fluor labeled chicken anti IgG (H+L), which was cross-adsorbed with a secondary antibody in 10% chicken serum (Invitrogen, Pleasanton, CA).

**Cilia imaging**. Human ARL13B was cloned using the In-Fusion HD Cloning system for seamless DNA cloning (Takara Bio USA, CA). The full-length ARL13B cDNA (428AA, NM_001174150) was cloned into pRRLSIN_cPP_PGKGFP_WPRE. The primers used for RT-PCR amplification of ARL13B werere: Fragment 1.FOR ctccccagggggatcatgttcagtctgatggccagttg, Fragment 1.REV ctcaccattgagatcacatcatgagcatcactgt, Fragment 2.FOR gatctcaatggtgagcaagggcgag, Fragment 2.REV ctctagaattacttgtacagctcgtccatgcc, Fragment 3.FOR caagtaattctagagtcgaccctgtggaatg, Fragment 3.REV gaggttgattgtcgatcaggcaccgggcttg.

***Lentivirus production and transduction***. 20 μg of a lentiviral vector, 15 μg of psPAX2 and 7.5 μg pMD2.G packaging mix were transfected into 293FT cells using the calcium phosphate transfection method in a 10-cm dish. The culture was incubated for 16 hours, and the medium was the replaced 24 h after transfection. Viral supernatants were harvested 48 and 72 hours after transfection, concentrated using the Lenti-X™ Concentrator (Takara Bio), and then stored at -80 °C. Single cells that were isolated from a hepatic organoid were incubated with virus (MOI 5-10) for 6 hours, and were then re-plated to allow the organoids to regenerate. ARL13B-GFP cilia within HOs were measured 7 days later with additional 2 days starvation.

***Single cell mass cytometry (CyTOF).*** Single cell preparations from hepatic organoids were generated using the methods described for scRNA-seq analysis. Then, the cell preparations were fixed with 2% paraformaldehyde at room temperature for 20 min, and then washed twice with PBS containing 0.5% BSA. The formaldehyde-fixed cells were incubated with metal-conjugated antibodies that were reactive with cell surface antigens

(**Table S5**) for 1 hour, washed once with PBS containing 0.5% BSA. The cells were then permeabilized with methanol for 15 minutes at 4 $^{0}$C, washed twice with PBS containing 0.5% BSA, and then incubated with metal-conjugated antibodies against intracellular antigens for 1 hour. The cells were then washed with PBS containing 0.5% BSA, and incubated at room temperature for 20 min with an iridium-containing DNA intercalator (Fluidigm) in PBS containing 2% paraformaldehyde. After intercalation/fixation, the cell samples were washed once with PBS containing 0.5% BSA, and twice with water before measurements were made using a CyTOF mass cytometer (Fluidigm). Normalization and de-barcoding were performed using previously described Matlab programs [63, 64]. After measurement and normalization, the individual data files were analyzed by first gating out doublets, debris and dead cells based on cell length, DNA content and cisplatin staining. tSNE maps were generated with software tools CYT, which is a graphical software package obtained from Dana Pe'er.

***RT-PCR analyses.*** Total RNA was extracted using TRIzol® RNA Isolation Reagents (Ambion, Grand Island, NY), and 2 µg RNA was reverse-transcribed using the iScript™ Advanced cDNA Synthesis Kit (BIO-RAD, Hercules, CA) according to the manufacturer's guidelines. The following TaqMan primer sets (Life Technologies Grand Island, NY) were used for the analyses: COL1A1 Hs00164004_m1, PDGFRB Hs01019589_m1, VIMENTIN Hs00958111_m1, ACTA2 (SMA), Hs00426835_g1, CDH1 Hs01023895_m1, KRT19 Hs00761767_s1, GAPDH Hs02786624_g1.

***Image Segmentation and Quantification****.* Immunostained and trichrome stained image processing and analysis was performed using the Fiji implementation of ImageJ. Quantification of the area of positive staining was calculated using the 'Trainable Weka Segmentation' plugin [65]. For training purposes, each measurement in ~5 to 10 areas of each analyzed element in every image (total 5 images) were used to train the classifiers by manually labelling all of the positive spots. Then, for experimental image analysis, the saved classifier was used to generate the probability map for each image. Target channels were then isolated, thresholded and binarized. For each treatment group, the

‘*Area Fraction*’ measured in 4 to 6 images were assessed. An unpaired Student’s t-test was used to test whether the measurements were significantly different.

For the quantification of collagen volume in 3-dimensional whole mount organoids that were stained with anti-COL1A antibodies, split channels including > 100 stack of images from 100 to 200 μm segments were used to calculate the volume. This was accomplished by multiplying the sum of the area of in each stack by the depth (volume = total area x depth). The collagen volume in measured in 4 organoids of each type were used to compare the collagen volume difference resulting from treatment with each drug. The macro script that we used for this analysis is available upon request.

***Single Organoid PCR***. Single organoids were isolated and transferred to 0.2 ml PCR tube containing mixture of 2.3 ul RLT lysis buffer (Qiagen), 1 ul dNTP and 1 ul Oligo-dT30VN (5′–AAGCAGTGGTATCAACGCAGAGTACT30VN-3′) (10 μM). Cell lysis was performed by incubating the samples at 72 °C for 3 min, and the tubes were immediately placed on ice. Full length of cDNA was generated by reverse transcription and terminal transferase (Smart-seq) in the RT mix, which included *SuperScript II reverse transcriptase, RNAse inhibitor, Superscript II first-strand buffe, DTT, Betaine, MgCl2, and TSO (5'AAGCAGTGGTATCAACGCAGAGTACATrGrG+G-3').* The resulting cDNA was then diluted 1:10, and the sample was used as the template for a TaqMan assay.

**Single organoid proline and hydroxyproline quantitation**. Individual organoids were formed in Nunclon™ Sphera™ 96 well microplates by seeding 5000 hepatoblast cells per well. The plates were then centrifuged at 500xg for 5 minutes. The drugs tested were: PDGFRB tyrosine kinase inhibitors (Crenolanib, Sunitinib and Imatinib); and a NOTCH inhibitor (DAPT). All anti-fibrotic reagents purchased from Selleckchem (TX, USA) or Tocris (MN, USA). The indicated concentration of each drug (10, 2, or 0.5 μM) was added to the microwell on day 12, and the drug was present for the last 10 days of organoid culture. The individual organoids were isolated and transferred to a 0.2 ml PCR tube containing 30 μl 6N HCL; and hydrolysis was performed at 120 °C for 100 minutes. The clear supernatant was transferred for further dilution and LC-MS analysis. Hydrolyzed organoid samples were brought to a 100 uL volume by addition dd$H_2O$. A 10 uL aliquot from each sample was mixed with an equal volume of an internal standard solution and then dried using a Speedvac (Thermo Fisher). Trans-4-Hydroxy-L-Proline (2,5,5-D3) and D7-L proline (Cambridge Isotope Laboratories, MA) were used as internal standards. The dried mixture was re-suspended in 50 uL water and derivatized with dansyl chloride (Sigma) using our previously describe modification [66-68] of the method developed by Guo and Lee [69]. LCMS analysis was then performed using an Agilent QTOF 6545 (Agilent, Santa Clara) coupled with a 1290 infinity I UHPLC (Agilent, Santa Clara). The samples were run on a Phenomenex C18 kinetic column. A mobile phase was 0.1% formic acid in water and B was 100% acetonitrile. The acquired data were analyzed using Masshunter Quantitative analysis software and the proline and 4-hydroxyproline concentrations were calculated based on a 10-point calibration curve.

*Data availability.* All raw single cell RNA-seq data and processed data have been deposited in the Gene Expression Omnibus (GEO) under accession GSE154883. Additional single cell RNA-seq dataset using for differentiation validation is obtained from our previous work at GSE139382. R scripts and macro enabling the main steps of the analysis are available from the corresponding authors upon request.

**Supplemental References**


1. Campagnola, P.J. & Loew, L.M. Second-harmonic imaging microscopy for visualizing biomolecular arrays in cells, tissues and organisms. *Nat Biotechnol* **21**, 1356-1360 (2003).
2. Deniset-Besseau, A. *et al.* Measurement of the second-order hyperpolarizability of the collagen triple helix and determination of its physical origin. *J Phys Chem B* **113**, 13437-13445 (2009).
3. Yamamoto, S. *et al.* Quantitative imaging of fibrotic and morphological changes in liver of non-alcoholic steatohepatitis (NASH) model mice by second harmonic generation (SHG) and auto-fluorescence (AF) imaging using two-photon excitation microscopy (TPEM). *Biochem Biophys Rep* **8**, 277-283 (2016).
4. Sun, W. *et al.* Nonlinear optical microscopy: use of second harmonic generation and two-photon microscopy for automated quantitative liver fibrosis studies. *J Biomed Opt* **13**, 064010 (2008).
5. Gailhouste, L. *et al.* Fibrillar collagen scoring by second harmonic microscopy: a new tool in the assessment of liver fibrosis. *J Hepatol* **52**, 398-406 (2010).
6. Drifka, C.R. *et al.* Comparison of Picrosirius Red Staining With Second Harmonic Generation Imaging for the Quantification of Clinically Relevant Collagen Fiber Features in Histopathology Samples. *J Histochem Cytochem* **64**, 519-529 (2016).
7. Trapnell, C. *et al.* The dynamics and regulators of cell fate decisions are revealed by pseudotemporal ordering of single cells. *Nat Biotechnol* **32**, 381-386 (2014).
8. Guan, Y. *et al.* Human Hepatic Organoids for the Analysis of Human Genetic Diseases. *JCI Insight* **2**, pii: 94954 (2017).
9. Guan, Y. *et al.* The phosphatidylethanolamine biosynthesis pathway provides a new target for cancer chemotherapy. *J Hepatol* **72**, 746-760 (2019).
10. Subramanian, A. *et al.* Gene set enrichment analysis: a knowledge-based approach for interpreting genome-wide expression profiles. *Proc Natl Acad Sci U S A* **102**, 15545-15550 (2005).
11. Verstockt, B. *et al.* Expression Levels of 4 Genes in Colon Tissue Might be Used to Predict Which Patients Will Enter Endoscopic Remission After Vedolizumab Therapy for Inflammatory Bowel Diseases. *Clin Gastroenterol Hepatol* (2019).
12. Wang, Z. *et al.* Identification of seven-gene signature for prediction of lung squamous cell carcinoma. *Onco Targets Ther* **12**, 5979-5988 (2019).
13. Labrecque, M.P. *et al.* Molecular profiling stratifies diverse phenotypes of treatment-refractory metastatic castration-resistant prostate cancer. *J Clin Invest* **130** (2019).
14. Merlos-Suarez, A. *et al.* The intestinal stem cell signature identifies colorectal cancer stem cells and predicts disease relapse. *Cell Stem Cell* **8**, 511-524 (2011).
15. Corominas-Faja, B. *et al.* Stem cell-like ALDH(bright) cellular states in EGFR-mutant non-small cell lung cancer: a novel mechanism of acquired resistance to erlotinib targetable with the natural polyphenol silibinin. *Cell Cycle* **12**, 3390-3404 (2013).
16. Davis, S. & Meltzer, P.S. GEOquery: a bridge between the Gene Expression Omnibus (GEO) and BioConductor. *Bioinformatics* **23**, 1846-1847 (2007).

17. Younossi, Z.M. *et al.* Global epidemiology of nonalcoholic fatty liver disease-Meta-analytic assessment of prevalence, incidence, and outcomes. *Hepatology* **64**, 73-84 (2016).
18. Sayiner, M., Koenig, A., Henry, L. & Younossi, Z.M. Epidemiology of Nonalcoholic Fatty Liver Disease and Nonalcoholic Steatohepatitis in the United States and the Rest of the World. *Clin Liver Dis* **20**, 205-214 (2016).
19. Tsuchida, T. & Friedman, S.L. Mechanisms of hepatic stellate cell activation. *Nature reviews. Gastroenterology & hepatology* **14**, 397-411 (2017).
20. Sircana, A., Paschetta, E., Saba, F., Molinaro, F. & Musso, G. Recent Insight into the Role of Fibrosis in Nonalcoholic Steatohepatitis-Related Hepatocellular Carcinoma. *International journal of molecular sciences* **20** (2019).
21. Marcher, A.B. *et al.* Transcriptional regulation of Hepatic Stellate Cell activation in NASH. *Sci Rep* **9**, 2324 (2019).
22. Tanaka, N. *et al.* Current status, problems, and perspectives of non-alcoholic fatty liver disease research. *World J Gastroenterol* **25**, 163-177 (2019).
23. Angulo, P. *et al.* Liver Fibrosis, but No Other Histologic Features, Is Associated With Long-term Outcomes of Patients With Nonalcoholic Fatty Liver Disease. *Gastroenterology* **149**, 389-397 e310 (2015).
24. Ramachandran, P. *et al.* Resolving the fibrotic niche of human liver cirrhosis at single-cell level. *Nature* **575**, 512-518 (2019).
25. Sato, Y. *et al.* Cholangiocytes with mesenchymal features contribute to progressive hepatic fibrosis of the polycystic kidney rat. *Am J Pathol* **171**, 1859-1871 (2007).
26. Moser, M. *et al.* A mouse model for cystic biliary dysgenesis in autosomal recessive polycystic kidney disease (ARPKD). *Hepatology* **41**, 1113-1121 (2005).
27. Kaimori, J.Y. *et al.* NEDD4-family E3 ligase dysfunction due to PKHD1/Pkhd1 defects suggests a mechanistic model for ARPKD pathobiology. *Sci Rep* **7**, 7733 (2017).
28. Perng, D.W. *et al.* Matrix metalloprotease-9 induces transforming growth factorbeta(1) production in airway epithelium via activation of epidermal growth factor receptors. *Life sciences* **89**, 204-212 (1989).
29. Kobayashi, T. *et al.* Matrix metalloproteinase-9 activates TGF-beta and stimulates fibroblast contraction of collagen gels. *Am J Physiol Lung Cell Mol Physiol* **306**, L1006-1015 (2014).
30. Talbot, J.J. *et al.* Polycystin-1 regulates STAT activity by a dual mechanism. *Proc Natl Acad Sci U S A* **108**, 7985-7990 (2011).
31. Ying, H.Z. *et al.* PDGF signaling pathway in hepatic fibrosis pathogenesis and therapeutics (Review). *Mol Med Rep* **16**, 7879-7889 (2017).
32. Vignais, M.L. & Gilman, M. Distinct mechanisms of activation of Stat1 and Stat3 by platelet-derived growth factor receptor in a cell-free system. *Mol Cell Biol* **19**, 3727-3735 (1999).
33. Bowman, T. *et al.* Stat3-mediated Myc expression is required for Src transformation and PDGF-induced mitogenesis. *Proc Natl Acad Sci U S A* **98**, 7319-7324 (2001).

34. Nevzorova, Y.A. *et al.* Overexpression of c-myc in hepatocytes promotes activation of hepatic stellate cells and facilitates the onset of liver fibrosis. *Biochim Biophys Acta* **1832**, 1765-1775 (2013).
35. Nicola, N.A. *et al.* Negative regulation of cytokine signaling by the SOCS proteins. *Cold Spring Harb Symp Quant Biol* **64**, 397-404 (1999).
36. Croker, B.A. *et al.* SOCS3 negatively regulates IL-6 signaling in vivo. *Nat Immunol* **4**, 540-545 (2003).
37. Liau, N.P.D. *et al.* The molecular basis of JAK/STAT inhibition by SOCS1. *Nature communications* **9**, 1558 (2018).
38. Rui, L., Yuan, M., Frantz, D., Shoelson, S. & White, M.F. SOCS-1 and SOCS-3 block insulin signaling by ubiquitin-mediated degradation of IRS1 and IRS2. *J Biol Chem* **277**, 42394-42398 (2002).
39. Kershaw, N.J., Laktyushin, A., Nicola, N.A. & Babon, J.J. Reconstruction of an active SOCS3-based E3 ubiquitin ligase complex in vitro: identification of the active components and JAK2 and gp130 as substrates. *Growth Factors* **32**, 1-10 (2014).
40. Timmermann, A., Kuster, A., Kurth, I., Heinrich, P.C. & Muller-Newen, G. A functional role of the membrane-proximal extracellular domains of the signal transducer gp130 in heterodimerization with the leukemia inhibitory factor receptor. *Eur J Biochem* **269**, 2716-2726 (2002).
41. Albrengues, J. *et al.* LIF mediates proinvasive activation of stromal fibroblasts in cancer. *Cell reports* **7**, 1664-1678 (2014).
42. Luo, K. Signaling Cross Talk between TGF-beta/Smad and Other Signaling Pathways. *Cold Spring Harbor perspectives in biology* **9** (2017).
43. Chakraborty, D. *et al.* Activation of STAT3 integrates common profibrotic pathways to promote fibroblast activation and tissue fibrosis. *Nature communications* **8**, 1130 (2017).
44. Hata, A. & Chen, Y.G. TGF-beta Signaling from Receptors to Smads. *Cold Spring Harbor perspectives in biology* **8** (2016).
45. Chen, X. *et al.* Integration of external signaling pathways with the core transcriptional network in embryonic stem cells. *Cell* **133**, 1106-1117 (2008).
46. Hsu, P.D. *et al.* DNA targeting specificity of RNA-guided Cas9 nucleases. *Nat Biotechnol* **31**, 827-832 (2013).
47. Cheng, J.X. & Xie, X.S. Coherent anti-Stokes Raman scattering microscopy: instrumentation, theory, and applications. *J. Phys. Chem. B* **108**, 827-840 (2004).
48. Enejder, A., Brackmann, C. & Svedberg, F. Coherent Anti-Stokes Raman Scattering Microscopy of Cellular Lipid Storage. *IEEE J. Sel. Top. Quantum Electron* **16**, 506-515 (2010).
49. Brackmann, C. *et al.* Nonlinear microscopy of lipid storage and fibrosis in muscle and liver tissues of mice fed high-fat diets. *J Biomed Opt* **15**, 066008-066001-066010 (2010).
50. Lin, J. *et al.* Assessment of liver steatosis and fibrosis in rats using integrated coherent anti-Stokes Raman scattering and multiphoton imaging technique. *J Biomed Opt* **16**, 116024 (2011).
51. Copple, B.L., Bai, S., Burgoon, L.D. & Moon, J.O. Hypoxia-inducible factor-1alpha regulates the expression of genes in hypoxic hepatic stellate cells important for collagen

deposition and angiogenesis. *Liver international : official journal of the International Association for the Study of the Liver* **31**, 230-244 (2011).
52. Rasband, W.S. (U.S. NIH, Bethesda, MD; 2019).
53. Hildebrand, T. & Ruegsegger, P. A new method for the model-independent assessment of thickness in three-dimensional images. *J. of Microscopy* **185**, 67-75 (1996).
54. Kang, H.M. *et al.* Multiplexed droplet single-cell RNA-sequencing using natural genetic variation. *Nat Biotechnol* **36**, 89-94 (2018).
55. Butler, A., Hoffman, P., Smibert, P., Papalexi, E. & Satija, R. Integrating single-cell transcriptomic data across different conditions, technologies, and species. *Nat Biotechnol* **36**, 411-420 (2018).
56. Kanehisa, M. & Goto, S. KEGG: kyoto encyclopedia of genes and genomes. *Nucleic Acids Res* **28**, 27-30 (2000).
57. Fabregat, A. *et al.* The Reactome pathway Knowledgebase. *Nucleic Acids Res* **44**, D481-487 (2016).
58. Stuart, T. *et al.* Comprehensive Integration of Single-Cell Data. *Cell* **177**, 1888-1902 e1821 (2019).
59. Wolf, F.A. *et al.* PAGA: graph abstraction reconciles clustering with trajectory inference through a topology preserving map of single cells. *Genome Biol* **20**, 59 (2019).
60. Wurmbach, E. *et al.* Genome-wide molecular profiles of HCV-induced dysplasia and hepatocellular carcinoma. *Hepatology* **45**, 938-947 (2007).
61. Lefebvre, P. *et al.* Interspecies NASH disease activity whole-genome profiling identifies a fibrogenic role of PPARalpha-regulated dermatopontin. *JCI Insight* **2** (2017).
62. Suppli, M.P. *et al.* Hepatic transcriptome signatures in patients with varying degrees of nonalcoholic fatty liver disease compared with healthy normal-weight individuals. *Am J Physiol Gastrointest Liver Physiol* **316**, G462-G472 (2019).
63. Zunder, E.R. *et al.* Palladium-based mass tag cell barcoding with a doublet-filtering scheme and single-cell deconvolution algorithm. *Nat Protoc* **10**, 316-333 (2015).
64. Finck, R. *et al.* Normalization of mass cytometry data with bead standards. *Cytometry A* **83**, 483-494 (2013).
65. Arganda-Carreras, I. *et al.* Trainable Weka Segmentation: a machine learning tool for microscopy pixel classification. *Bioinformatics* **33**, 2424-2426 (2017).
66. Wu, M. *et al.* Opiate-induced Changes in Brain Adenosine Levels and Narcotic Drug Responses. *Neuroscience* **228**, 235-242 (2013).
67. Park, W. *et al.* Metabolomic Markers Differentiate Mucinous and Non-Mucinous Pancreatic Cysts. *Gastrointestinal Endoscopy* **78**, 295-302 (2013).
68. Manhong Wu *et al.* Increased Dipeptide Abundance in Non-Small Cell Lung Cancer *Rapid Commun. Mass Spectrom.* **27**, 2091-2098 (2013).
69. Guo, K. & Li, L. Differential 12C-/13C-isotope dansylation labeling and fast liquid chromatography/mass spectrometry for absolute and relative quantification of the metabolome. *Anal Chem* **81**, 3919-3932 (2009).
70. Aubrey, B.J., Kelly, G.L., Janic, A., Herold, M.J. & Strasser, A. How does p53 induce apoptosis and how does this relate to p53-mediated tumour suppression? *Cell Death Differ* **25**, 104-113 (2018).

71. Ou, D.L. *et al.* Induction of DNA damage-inducible gene GADD45beta contributes to sorafenib-induced apoptosis in hepatocellular carcinoma cells. *Cancer Res* **70**, 9309-9318 (2010).

**Table S1**. Classification of the cell types found within the 15 clusters identified in human hepatic organoids. The transcriptomes of the organoid cell clusters are compared with that of non-hepatocyte cells present in control and cirrhotic human livers [24]. The comparisons were performed using the Seurat label transfer function [55]. Based upon the highest level of concordance between the clusters and the sequences present in the previously defined cell types identified in human liver: Clusters 0, 1, 3, 4, 6 and 7 are identified as scar-associated mesenchymal cells (**SAMe**); cluster 2 as mesothelia; cluster 10 as cholangiocytes; and cluster 14 as endothelia. Since the reference sequences only include a very limited number of hepatocytes, cell clusters containing hepatocytes could not be identified by this analysis. However, because mRNAs (*ALB, TTR, HNF4A and SERPINA1*) encoding known hepatocyte markers were expressed in cluster 5, they were identified as hepatocyte precursor cells (see Fig. S4)). Although a high proportion of cluster 8 and 9 cells were labeled as endothelial cells by the Seurat analysis, because the Pearson analysis indicates that their correlation with reference endothelial cells is ~0.5, they were identified as early endothelial cells. The identities of clusters 11, 12 and 13 could not be assigned with certainty. Numbers within the parentheses located next to cell type indicators correspond with the subpopulations annotated in [24].

| **Cluster:** | 0 | 1 | 2 | 3 | 4 | 5 | 6 | 7 | 8 | 9 | 10 | 11 | 12 | 13 | 14 |
|---|---|---|---|---|---|---|---|---|---|---|---|---|---|---|---|
| Cholangiocyte(2) | 0.00 | 0.00 | 0.01 | 0.00 | 0.01 | 0.50 | 0.00 | 0.00 | 0.00 | 0.00 | 0.11 | 0.00 | 0.00 | 0.02 | 0.00 |
| Cholangiocyte(3) | 0.00 | 0.00 | 0.00 | 0.00 | 0.00 | 0.42 | 0.00 | 0.00 | 0.00 | 0.00 | **0.57** | 0.00 | 0.01 | 0.04 | 0.00 |
| Endothelia (5) | 0.00 | 0.00 | 0.00 | 0.00 | 0.00 | 0.00 | 0.00 | 0.00 | 0.00 | 0.00 | 0.00 | 0.00 | 0.00 | 0.00 | 0.32 |
| Endothelia (6) | 0.00 | 0.00 | 0.00 | 0.00 | 0.02 | 0.00 | 0.02 | 0.00 | 0.00 | 0.00 | 0.00 | 0.17 | 0.03 | 0.19 | **0.41** |
| Endothelia (7) | 0.00 | 0.00 | 0.00 | 0.00 | 0.02 | 0.01 | 0.02 | 0.00 | **0.80** | **0.93** | 0.09 | 0.14 | 0.06 | 0.57 | 0.21 |
| Mesothelia | 0.00 | 0.02 | **0.57** | 0.00 | 0.06 | 0.01 | 0.10 | 0.03 | 0.00 | 0.01 | 0.02 | 0.00 | 0.00 | 0.00 | 0.00 |
| MPs (8) | 0.00 | 0.00 | 0.00 | 0.00 | 0.00 | 0.00 | 0.00 | 0.00 | 0.00 | 0.00 | 0.00 | 0.00 | 0.02 | 0.00 | 0.00 |
| SAMe | **0.96** | **0.77** | 0.38 | **1.00** | **0.83** | 0.03 | **0.85** | **0.97** | 0.20 | 0.05 | 0.20 | 0.52 | 0.28 | 0.04 | 0.06 |
| Tcells (1) | 0.02 | 0.11 | 0.03 | 0.00 | 0.04 | 0.03 | 0.01 | 0.00 | 0.00 | 0.00 | 0.00 | 0.15 | 0.57 | 0.13 | 0.00 |
| Tcells (2) | 0.01 | 0.09 | 0.02 | 0.00 | 0.01 | 0.00 | 0.00 | 0.00 | 0.00 | 0.00 | 0.00 | 0.02 | 0.04 | 0.01 | 0.00 |

| Gene | p_val | FC | % ARPKD | % Control | p_val_adj |
|---|---|---|---|---|---|
| *PCSK1N* | 5.09E-53 | 7.65 | 0.84 | 0.16 | 1.22E-48 |
| *MT2A* | 7.63E-31 | 7.23 | 0.92 | 0.57 | 1.84E-26 |
| *THY1* | 5.66E-41 | 6.76 | 0.66 | 0.07 | 1.36E-36 |
| *KRT17* | 8.8E-26 | 5.32 | 0.82 | 0.46 | 2.12E-21 |
| *COL1A2* | 1.29E-31 | 5.19 | 0.90 | 0.50 | 3.1E-27 |
| *SAA1* | 5.19E-48 | 4.83 | 0.95 | 0.47 | 1.25E-43 |
| *TGFBI* | 3.05E-32 | 4.78 | 0.89 | 0.52 | 7.35E-28 |
| *CCDC80* | 2.4E-39 | 4.76 | 0.87 | 0.33 | 5.77E-35 |
| *VIM* | 4.05E-39 | 4.31 | 0.98 | 0.59 | 9.76E-35 |
| *COL1A1* | 2.02E-19 | 4.19 | 0.83 | 0.56 | 4.85E-15 |

**Table S2.** The 10 most differentially expressed genes identified by comparing the transcriptomes of cholangiocytes (cluster 10) in control and ARPKD organoids. This table shows the gene symbol; the fold change (FC), which is the ratio of the expression level in ARPKD relative to control cholangiocytes; the raw and Bonferroni-adjusted p-values for each comparison; and the percentage of cholangiocytes in ARPKD or control organoids that express the indicated mRNA. The Wilcoxon Rank Sum test was used to identify the differentially expressed genes for the two groups of cells.

**Table S3 is at the end of this file**

**Table S4**. The primary antibodies used for immunohistochemistry and their sources.

| Symbol | NAME | CLONE | VENDOR | CAT# |
|---|---|---|---|---|
| A1AT | α-1-Antitrypsin | | DAKO | A001202 |
| Acetylated Tubulin | mouse anti-acetylated tubulin, | 6-11B-1 | Sigma | T7451 |
| ALB | Human Albumin Antibody | | Bethyl | A80-129A |
| COL1A1 | Collagen1 | | Abcam | ab34710 |
| ECAD | Anti-E-Cadherin | 36 | BD | 610182 |
| EpCAM | Purified anti-human CD326 (EpCAM) Antibody | 9C4 | Biolegend | 324202 |
| HNF4A | HNF4a (C-19) | | Santa Cruz | sc6556 |
| HNF4A | HNF4A | | Abcam | ab199431 |
| Ki67 | Ki-67 Antibody (H-300) | | Santa Cruz | sc-15402 |
| KRT18 | CK18 | DC 10 | DAKO | M 7010 |
| KRT19 | Cytokeratin 19 | RCK108 | DAKO | M088801-2 |
| KRT19 | Cytokeratin 19 Antibody | A53-B/A2 | Santa Cruz | sc-6278 |
| KRT19 | Krt19 Antibody | TROMA-III | DSHB | TROMA-III |
| KRT7 | Cytokeratin 7 | | DAKO | M701801-2 |
| KRT8 | Anti-Cytokeratin 8 antibody | EP1628Y | Abcam | (ab53280) |
| KRT8 | Krt8 Antibody | TROMA-I | DSHB | TROMA-I |
| SMA | Actin, Smooth Muscle | 1A4 | Cell marque | 1A4 |
| SMA | Anti-alpha smooth muscle Actin | | Abcam | ab5694 |
| SMA | Monoclonal Anti-a Smooth Muscle Actin | 1A4 | Sigma | A2547 |
| SMA | α-Actin Antibody (1A4): | 1A4 | Santa Cruz | sc-32251 |
| SOX9 | Anti-Sox9 Antibody | | Millipore | AB5535 |
| SOX9 | Human SOX9 Antibody | | R&D | AF3075-SP |
| VANGL1 | Vang-like Protein 1/VANGL1 | | Novusbio | NBP1-86990 |
| ZO-1 | ZO-1 | ZO1-1A12 | Life | 339100 |
| ZO-1 | ZO-1 | | Life | 402200 |
| PDGFRB | Recombinant Anti-PDGFR beta antibody | Y92 | Abcam | (ab32570) |
| PDGFRB | Human PDGF R beta | | R&D | AF385 |
| PDGFRB | PDGF Receptor β | 28E1 | Cell Signaling Technology | 3169 |
| CD56 | NCAM | 123C3 | Invitrogen | 07-5603 |
| b-Catenn | Non-phospho (Active) β-Catenin | | Cell Signaling Technology | 8814 |
| JAG1 | Polyclonal Ab | | R&D | AF1277-SP |

| | | | | |
|---|---|---|---|---|
| JAG1 | Jagged1 Antibody (C-20) | | Santa Cruz | sc-6011 |
| JAG1 | JAG1 | TS1.15H | DSHB | TS1.15H |
| NOTCH1 | Anti-activated Notch1 | | Abcam | ab8925 |
| NOTCH1 | Notch1 intracellular domain (human) | bTAN 20 | DSHB | bTAN 20 |

**Table S5.** Antibodies used for CyTOF analyses.

| Metal Isotope | Mass | 1st | 2nd |
|---|---|---|---|
| Y | 89 | NA | CD45 |
| In | 113 | ALBUMIN | ALBUMIN |
| In | 115 | Vimentin | Vimentin |
| La | 139 | NA | COL1 |
| La | 140 | NA | CD31 |
| Pr | 141 | CD326 EpCAM | CD326 EpCAM |
| Nd | 142 | CD26 | CD19 |
| Nd | 143 | CD117 | CD117 |
| Nd | 144 | Tra1-81 | CD11c |
| Nd | 145 | ampk | Desmin |
| Nd | 146 | SOX17 | CD3 |
| Sm | 147 | CK7 | CK7 |
| Nd | 148 | FOXA2/HNF3b 148 | CD68 |
| Sm | 149 | Histone H3 | CD271 |
| Nd | 150 | AFP (3150025B) | AFP (3150025B) |
| Eu | 151 | CD29 ITGB1 | PGC1 |
| Sm | 152 | JAG1_G | JAG1_G |
| Eu | 153 | pSTAT3 | CD112 |
| Sm | 154 | GATA4 | KDR (CD309) |
| Gd | 155 | HNF4A | HNF4A |
| Gd | 156 | pSTAT3 | CD140B |
| Gd | 157 | Tra1-60 | ApoE |
| Gd | 158 | ECAD | CD169 |
| Tb | 159 | CD90 | CD11c |
| Gd | 160 | SOX2 | PDGFRa |
| Dy | 161 | LGR5 (3161025B) 161 | GFPT2 |
| Dy | 162 | CD49d | CD49d |
| Dy | 163 | pAkt(T308) | CD54 |
| Dy | 164 | cyclinB | Podoplanin |
| Ho | 165 | phospho Rb pS807/pS811 | CD163 |
| Er | 166 | SOX9 | ARG1 |
| Er | 167 | HNF1B | SMA_R |
| Er | 168 | Ki67 | CD206 |
| Tm | 169 | conexin43 | conexin43 |
| Er | 170 | CK19 _IN | CK19 _IN |
| Yb | 171 | CD49f | PDL1 |
| Yb | 172 | CD31 | CD31 |
| Yb | 173 | A1AT | A1AT |
| Yb | 174 | Keratin (CK8/18) (3174014A) | CD144 |
| Lu | 175 | CD184 | NR1D1 |
| Yb | 176 | CD56 NCAM | CD56 |
| Ir | 191/193 | DNA | DNA |
| Pt | 195 | cisplatin | Cisplatin |
| Bi | 209 | NA | CD47 |

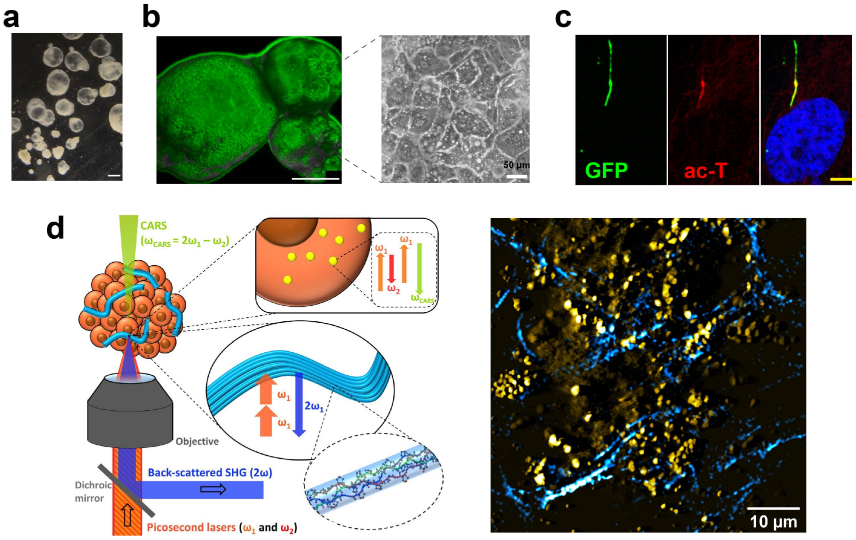


**Figure S1. Hepatic organoid culture system and SHG analysis. a**, A low power, bright field view of HOs obtained after 21 days of differentiation. Scale bar is 500 μm. **b**, **Left**, Calcein AM staining indicates that cells within an organoid are viable, Scale bar is 500 μm. **Right**: a high-power bright field image (of the region indicated in the middle image) shows the polygonal hepatocyte morphology of the cells within a HO. These cells also have lipid vesicles (bright areas). **c**, A subpopulation of cells within an HO have primary cilium. A primary cilium of a day 21 HO was visualized by expression of ARL13B-GFP fusion protein (GFP, green) and immunostaining with an anti-acetylated tubulin antibody (ac-T, red); cell nuclei were counter-stained with DAPI (blue) in the merged image (right). Scale bar is 5 μm. **d**, **Left**: Schematic of SHG and CARS microscopy of a HO with collagen fibers (cyan). Two excitation beams at frequencies $\omega_1$ and $\omega_2$ were focused on the sample by a high-numerical aperture (1.45, 100x) objective. The endogenous SHG signal, constructively built up at double the $2\omega_1$ frequency and emitted from the non-centrosymmetric collagen fibers (as shown in the inset), was collected in back-reflection mode. The CARS signal emitted at $\omega_{CARS} = 2\omega_1 - \omega_2$ by the intracellular lipid stores (as shown in the insert) was simultaneously collected in transmission mode. **Right**: A merged CARS/SHG image of a day 21 control hepatic organoid. The lipids are yellow, and the collagen fibers are cyan colored.  Scale bar, 10 μm.

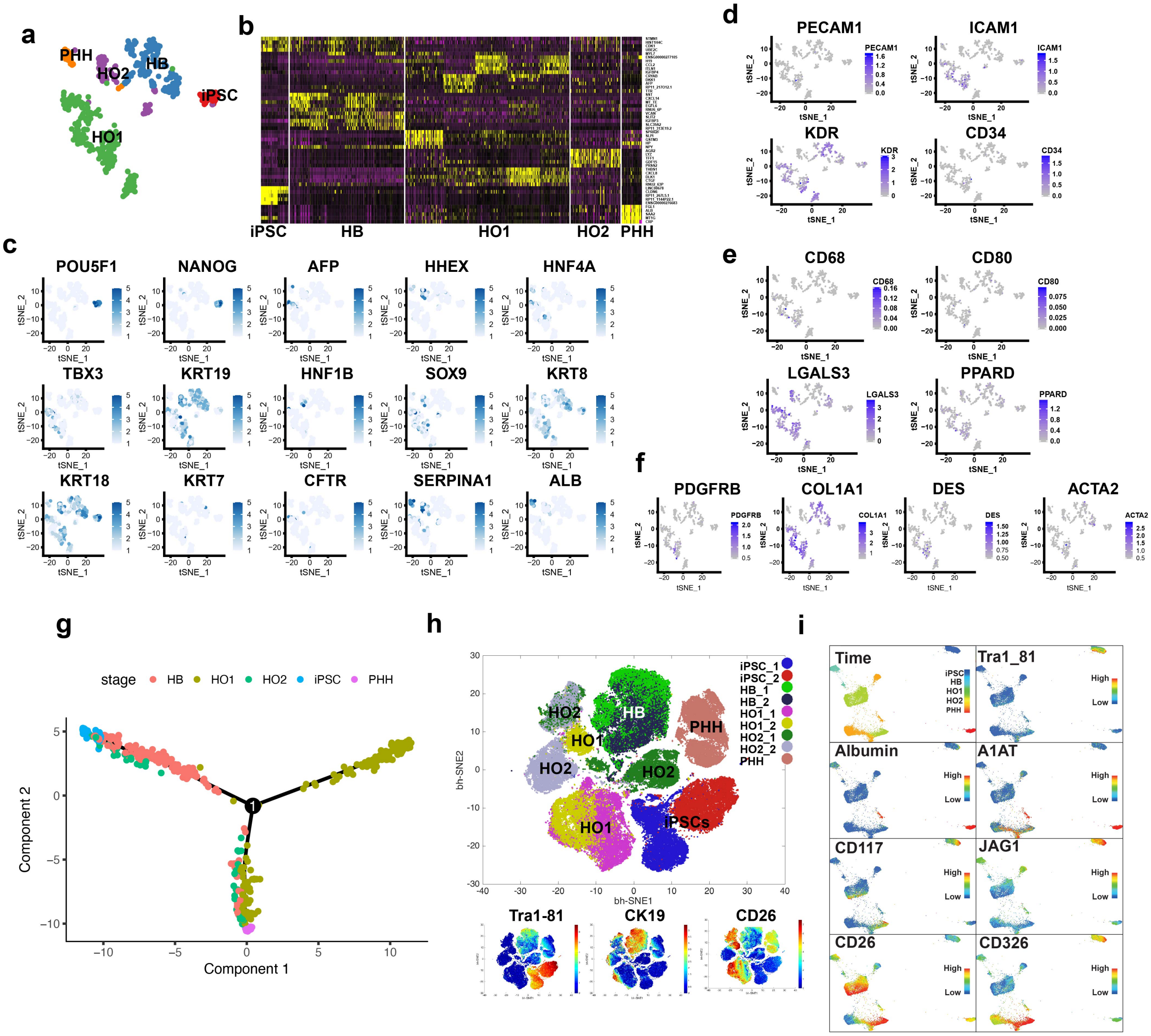


**Figure S2. Transcriptomic analysis reveals that hepatic organoids have cells of multiple lineages. a,** tSNE plot of scRNA-seq data generated from organoid cultures at five different stages of differentiation: iPSC, hepatoblasts (HB), and primary (HO1) and secondary (HO2) organoids. Primary human hepatocyte (PHH) data were used as the positive control for differentiation. The cells at each differentiation stage were separated into distinct clusters that are indicated by color. **b,** A heat map showing the marker genes for each of the five differentiation clusters; the top 5 (or fewer if <5 were found) marker genes are color-coded for each cluster. Representative genes for each cluster are shown on the right. Each column is an individual cell within the cluster indicated at the bottom; and each gene is in a row. A Wilcoxon rank sum test was used to test for differential expression of the mRNAs; all markers were expressed in >25% of the cells; and the threshold for selecting a marker was set at a minimum of $\log_2$ (fold-change) > 0.25. **c-f**, tSNE projection of scRNA-Seq data for 559 cells. The color of each cell (represented by a dot) is based on the normalized level of expression of canonical markers for: iPSC (PU5F1, NANOG), hepatoblasts (AFP, HHEX, HNF4A, TBX3, KRT19, KRT8), hepatocytes (SERPINA1, ALB, CK18), cholangiocytes (KRT7, CFTR, SOX9, HNF1B), endothelial cells (PECAM1, ICAM1, KDR, CD34), Kupffer cells (CD68, CD80, LGALS3, PPARD) or hepatic stellate cells (PDGFRB, COL1A, DES, ACTA2). **g**. A Monocole plot shows the pseudotime representation of the cellular differentiation trajectory. iPSC (shown on the left) through the hepatoblast (HB) stage, which then continue on to the various cell types present primary (HO1) and secondary (HO2) organoids and primary human hepatocytes (PHH). **h**, **Top**: A bhSNE map, which is generated using CyTOF data obtained with 38 antibodies, shows the clustering of cells generated from iPSC, hepatoblasts, HO1, HO2 and PHH. The cells are separated into spatially distinct subsets based on the combination of markers that they express. Each point in the bhSNE map represents an individual cell. **Bottom:** The cells in the bhSNE map are colored according to the intensity of expression of differentiation stage markers, which include: Tra1-81 for iPSC; CK19 for hepatoblasts; and CD26 for hepatocytes and cholangiocytes. **i.** A Force Directed Layout (FDL) map shows the differentiation trajectory along the iPSC-HB-HO1-HO2-PHH axis. Canonical markers - including Tra1-81, ALB, A1AT, CD26, JAG1, CD117 and CD326 - were projected on the FDL maps.

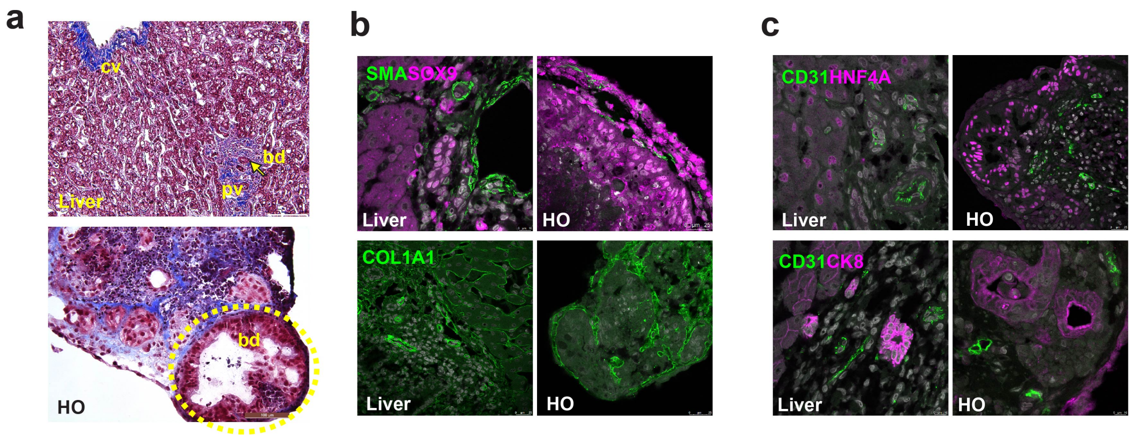


**Figure S3. HOs form complex structures that resemble those found in human liver**. **a,** Trichrome staining shows the similarity between some of the structures present in normal liver (Top) and HOs (bottom). The following structures are indicated in normal liver: cv, central vein; pv, portal vein; and bd, bile duct. The dotted circle indicates a ductal area in the hepatic organoid. **b,c**, HOs were immunostained with antibodies to different cell types found in liver: mesenchymal cells (SMA), endothelial cells (CD31), cholangiocytes (SOX9), and hepatocytes (HNF4A, CK8). Anti-CD31 staining indicates that structures resembling vessels are formed in HOs. The anti-SMA and anti-COL1A1 staining identifies collagen and mesenchymal cells, which are found in peri-ductal and vascular areas.

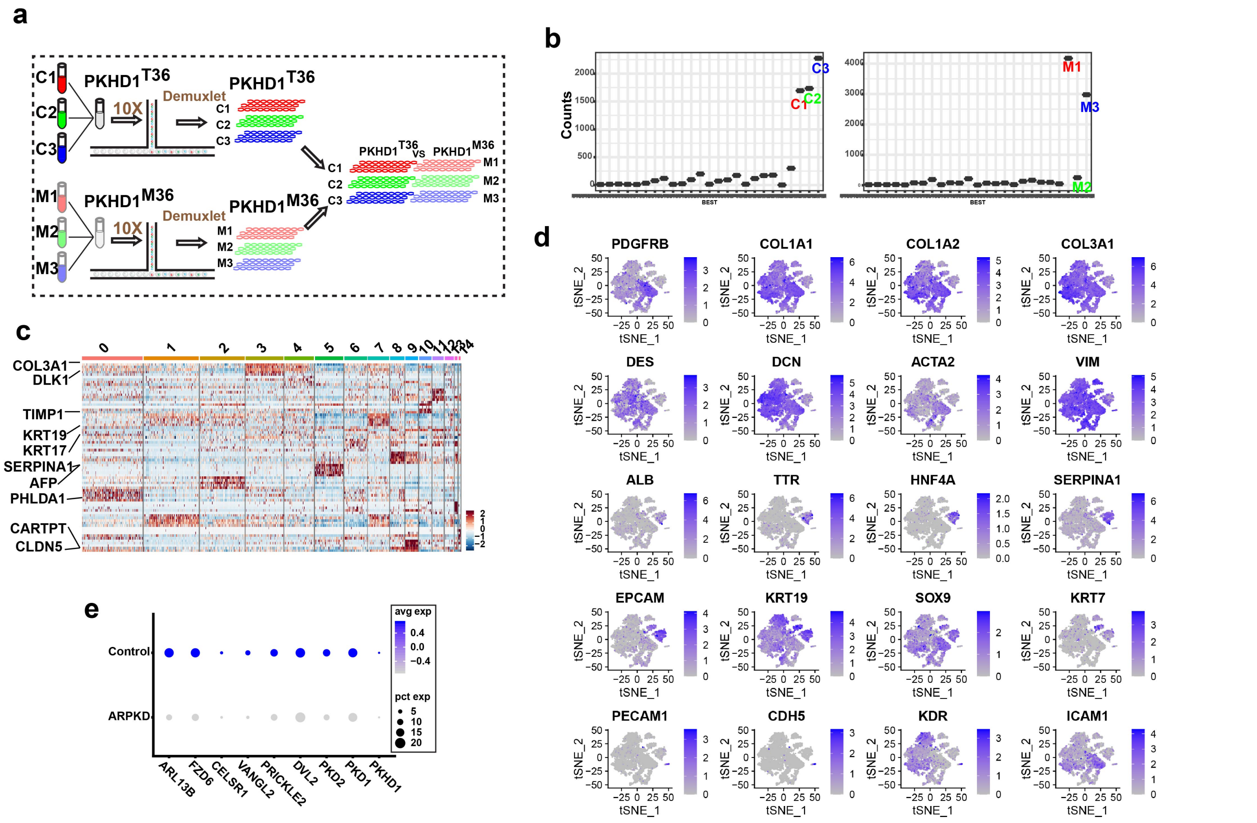


**Figure S4**. **scRNA-seq analysis HO differentiation**. **a**, The pipeline for the multiplex analysis of the scRNA-Seq data generated from control HOs prepared from 3 unrelated individuals (C1-C3), and from ARPKD organoids that were prepared from isogenic iPSCs (M1, M2, M3) with an engineered *PKHD1 M36* mutation. Pooled samples of control and ARPKD organoids were separately analyzed, and the multiplexed data was deconvoluted based upon analysis of allelic differences using 'demuxlet' software [54]. **b**, Dot plots show a summary the number of cells identified for each subject by 'demuxlet' in control (left) and ARPKD (right) pooled samples. Dots that are close to base line represent doublet or ambiguous droplets that were not analyzed. **c,** A heat map showing marker genes for the 15 clusters. Each cluster is represented by the top 5 marker genes, which are color-coded for each cluster; and representative genes for each cluster are shown on the left. Each column is an individual cell within the indicated cluster at the top; and each gene is in a row. A Wilcoxon rank sum test was used to test for differential expression of the mRNAs; all markers were expressed in >25% of the cells; and the threshold for selecting a marker was set at a minimum of $\log_2$ (fold-change) > 0.25. **d**, The level of expression of mRNAs for known cell type-specific markers are visualized by projection on the t-SNE map. These plots show the level of expression of mRNAs that encode known markers for: mesenchymal cells and myofibroblasts (*PDGFRB, COL1A1, COL1A2, COL3A1, DES, DCN,*

*ACTA2* and *VIM*); hepatocytes (*ALB, TTR, HNF4A* and *SERPINA1*); cholangiocytes (*EPCAM, KRT19, SOX9* and *KRT7*); and endothelial cells (*EPCAM1, CDH5, KDR* and *ICAM1*). **e**, The levels of expression of mRNAs associated with the primary cilium and planar cell polarity (PCP) are decreased in ARPKD hepatic organoids. Based upon analysis of scRNA-Seq data for all cells within an organoid, the dot size indicates the percentage of cells expressing the indicated mRNA. The dot color indicates the average level of expression of the indicated mRNA: blue indicates a high, while grey indicates a low level of expression. The level of expression of multiple mRNAs associated with PCP (*FZD6, CELSR1, VANGL2, PRICKLE2, DVL2*) or primary cilium (*ARL13B, PKD2, PKD1, PKDH1)* are decreased in ARPKD organoids relative to their isogenic control organoids; while the percentages of cells expressing each of these mRNAs are not altered.

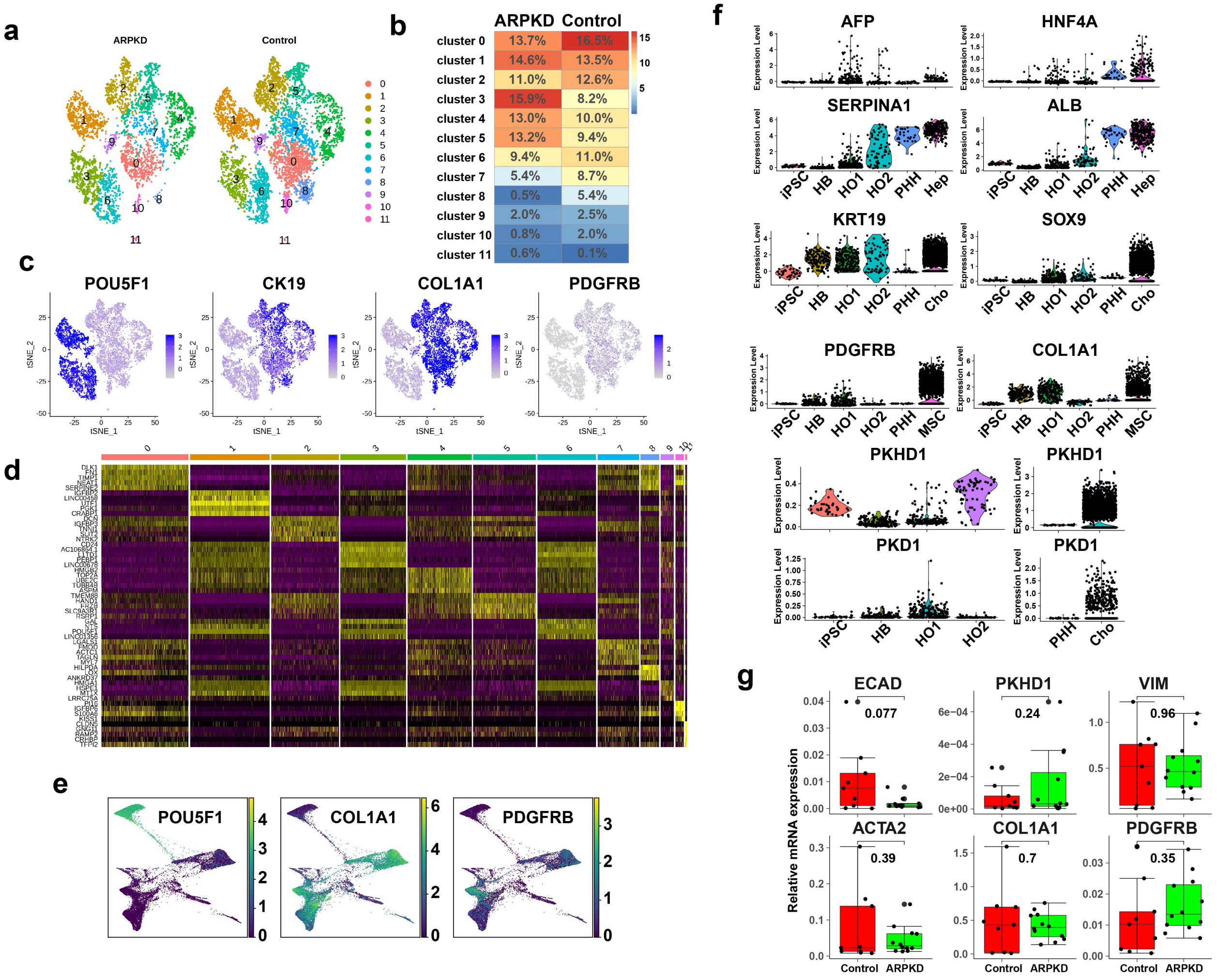


**Figure S5. scRNA-seq analysis of iPSC and hepatoblasts (HB)**. scRNA-Seq data was obtained from ARPKD and isogenic control cells at the HB (day 9) and iPSC (day 0) stages prepared from 3 different donors. **a**, t-SNE embeddings identify 12 cell clusters (clusters 0-11) that are indicated by a different color. **b**, This heatmap shows the percentage of cells within each cluster for the ARPKD and their isogenic control cells; and the box color represents the percentage of cells in that cluster according to the indicated scale. **c**, The levels of expression of 4 mRNAs were visualized by projection on the t-SNE map. *POU5F1 (OCT4)* mRNA is only expressed in iPSCs; while *COL1A1, KRT19*, and *PDGFRB* mRNAs are expressed in HBs. **d**, A heat map showing the relative level of expression of 5 selected mRNAs for each of the 12 clusters shown in **a**. The gene symbols for the selected markers are shown on the left; and groups of 5 genes are indicated from top to bottom for cluster 0 through 11, respectively. For the analysis: a Wilcoxon rank sum test was used to test for differential mRNA expression; all markers had to be expressed in >25% of the cells; and the marker selection threshold was set at a minimum of log2 (fold-change) > 0.25. **e**, Canonical markers were projected on PAGA differentiation trajectory maps. **f**, Violin plots showing the level of expression of mRNAs encoding markers for hepatic progenitor cells (*AFP*), hepatocytes (*HNF4A, SERPINA1, ALB*), cholangiocytes (*KRT10, SOX9*), myofibroblasts (*PDGFRB, COL1A1*) and primary cilium (*PKDH1, PKD1*) during HO development. scRNA-Seq data was generated from iPSC, day 9 hepatoblast (HB), and in primary (HO1) and secondary (HO2) organoid cultures. For comparison purposes, scRNA-Seq from primary human hepatocyte (PHH), and published scRNA-Seq data for cholangiocytes (Cho) and mesenchymal cells (MSC) in human liver tissues was used for this analysis. The secondary organoid (HO2) cultures are formed by dissociating HO1 organoids into single cells, which will then reform organoids in 12 days. Each dot shows the level of expression of the indicated mRNA at the indicated stage as determined by analysis of the scRNA-Seq data. *PKDH1* and *PKD1* mRNAs are expressed at a lower level than other mRNAs in the organoid cultures; and because of this, their expression levels in human hepatocytes and in cholangiocytes in human liver are shown as separate graphs. Of note, the primary cilium mRNAs are expressed at the hepatoblast and HO stages in the organoid cultures, which means they are expressed during the period when hepatoblasts differentiate into hepatocytes and cholangiocytes. **g**, Cells in day 9 control and ARPKD organoid cultures have equivalent levels of expression of mRNAs encoding myofibroblast markers. RNA was prepared from day 9 (hepatoblast stage) differentiating control and ARPKD organoid cultures (n=3 donors, each measured in duplicate). RT-PCR was used to measure the level of expression of mRNAs encoding 5 genes that are expressed in myofibroblasts, and the level of *PKDH1* mRNA expression is also shown. Each dot shows the measurement obtained from one culture; the vertical line is the average expression level; and the box plot covers the 25 to 75 percentiles. The results of a 't-test' comparing the expression level for each mRNA in ARPKD and control cultures is also shown.

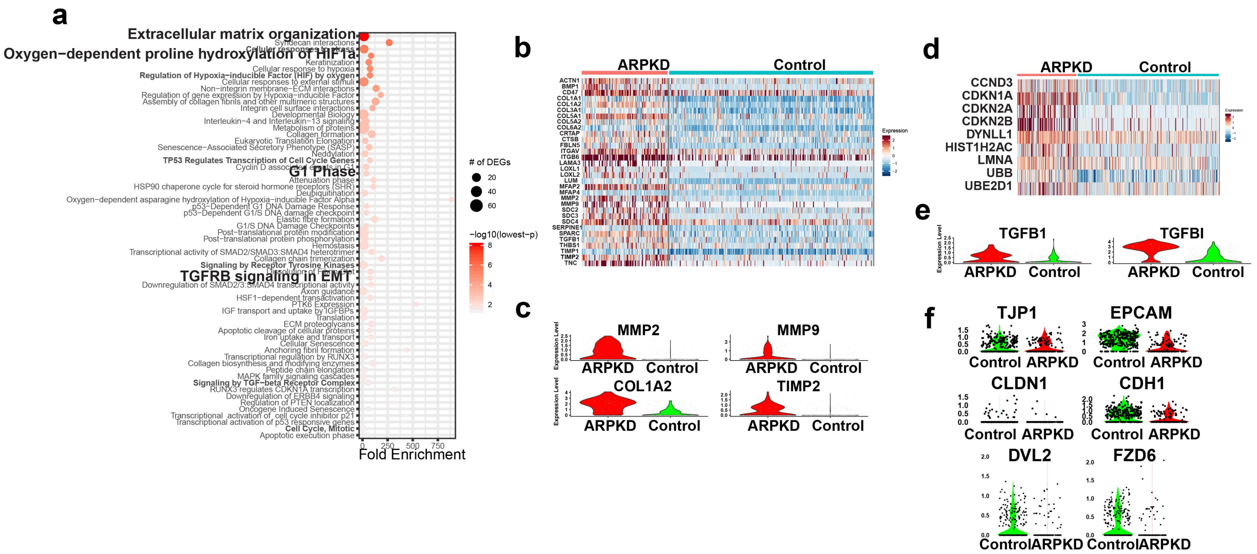


**Figure S6. Pathways activated in ARPKD cholangiocytes. a**, The level of expression of 439 mRNAs were altered in ARPKD cholangiocytes relative to control cholangiocytes (Fold Change > 1.3, minimum fraction of cells >0.1). Reactome Pathway Database (https://reactome.org/) analysis of the 439 differentially expressed identified 62 significantly enriched pathways. The size of the circle adjacent to each pathway indicates the number of differentially expressed genes (DEG) within that pathway; and the color indicates the $-\text{Log}_{10}$ of the p-value for the enrichment. The Extracellular Matrix organization (ECMO) pathway was most significantly enriched pathway; it had over 32 genes whose expression was up-regulated in ARPD cholangiocytes and an enrichment p-value of $8.4 \times 10^{-9}$. **b**, Heatmaps show ECMO pathway mRNAs with an increased level of expression in ARPKD cholangiocytes. **c**, Violin plots show the increased level of expression of ECMO mRNAs (*MMP2,* 2.6-fold increase*, MMP9, 1.7-fold; COL1A2,* 5.2-fold and*TIMP2,* 1.5-fold) that are increased in ARPKD cholangiocytes. **d**, Heatmaps show the genes within cell cycle and mitosis pathways have an increased level of expression in ARPKD cholangiocytes. **e**, Violin plots TGFβ signaling mRNAs (*TGFB1,* 1.3-fold*; TGFBI,* 4.8-fold*)* that are increased in ARPKD cholangiocytes. **f**, Violin plots show the level of expression of mRNAs that encode known markers for ductal epithelium and tight junction proteins (*CLDN1, CLDH1, TJDH1, EPCAM*) and planar cell polarity (PCP) (*DVL2, FZD6*) within cluster 10 cells in control and ARPKD hepatic organoids.

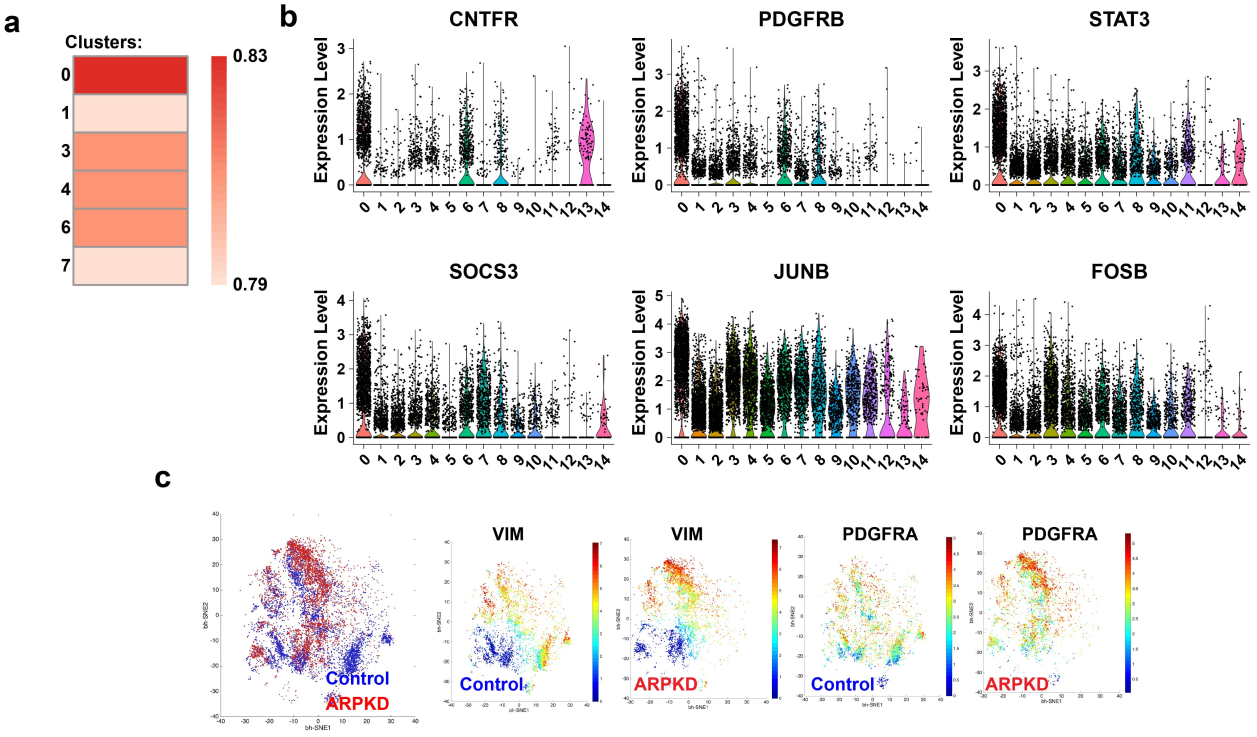


**Figure S7**. **Cluster 0 cells are myofibroblasts. a**, A heatmap showing the level of correlation between the transcriptomes of the 6 mesenchymal cell clusters (0, 1, 3, 4, 6, and 7) identified in hepatic organoids and myofibroblasts, which are the scar associated mesenchymal cells found in cirrhotic human liver tissue [24]. The extent of linear correlation (Pearson) between the 3000 variable features in each mesenchymal cluster with that of myofibroblasts is indicated in by the box color. The dark red box indicates that cluster 0 has the highest level of correlation with myofibroblasts. **b**, Violin plots show the level of 6 representative markers mRNAs that are expressed at higher levels in cluster 0 cells versus the 14 other clusters. **c**, ***Left panel***: A bhSNE map generated using CyTOF data from 40 antibodies shows the clustering of cells generated from control and ARPKD organoids. The cells are separated based on the combination of markers that they express, and are color coded according to whether they are from ARPKD or control organoids. Each point in the bhSNE map represents an individual cell. ***Right panels***: bhSNE maps are prepared for ARPKD or control organoids, and the individual cells are colored according to the intensity of VIM or PDGFRA protein expression.

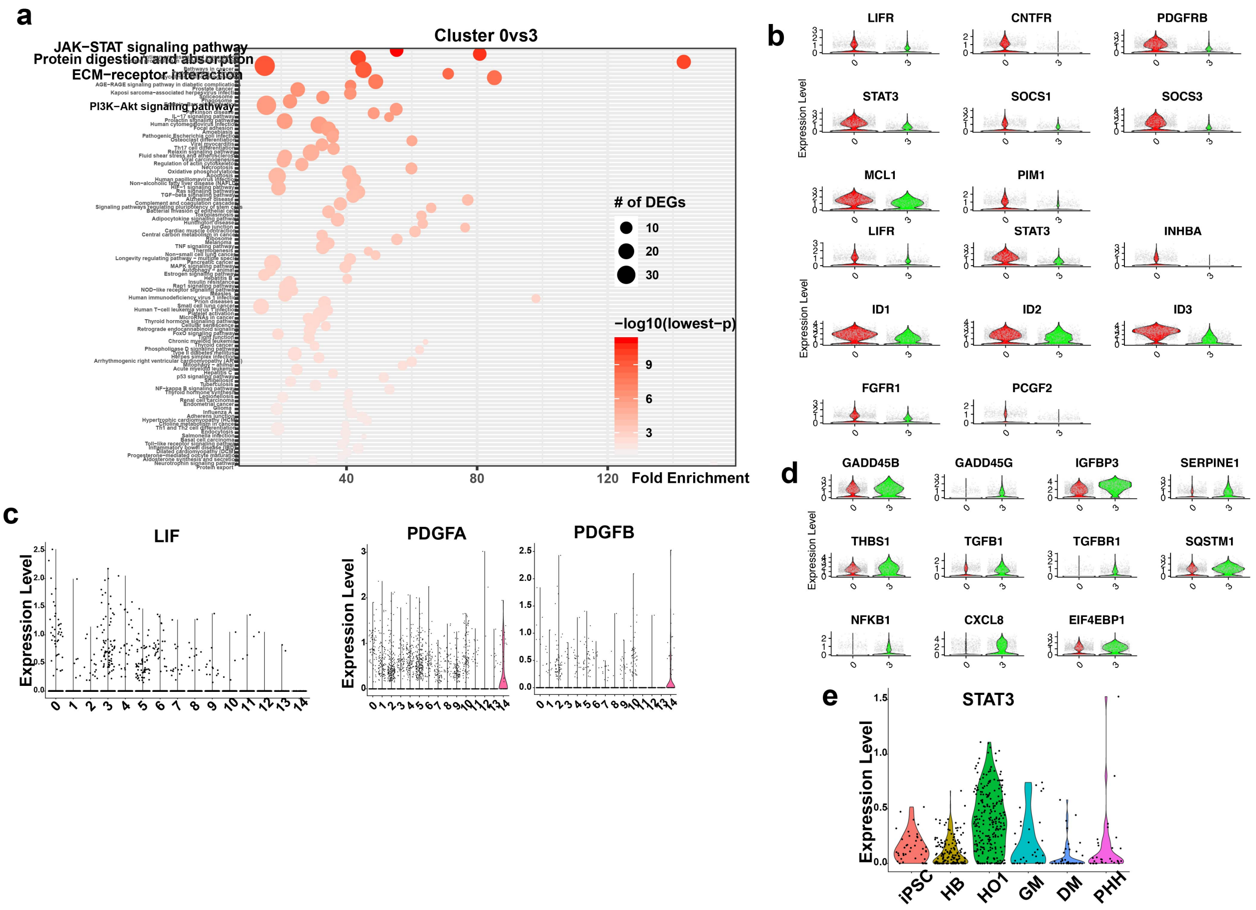


**Figure S8**. **Pathways activated in ARPKD myofibroblasts. a**, Analysis of 682 differentially expressed genes in cluster 0 (versus cluster 3) cells identified 106 significantly enriched pathways. Of these, the JAK-STAT signaling, ECM-receptor interaction, and protein digestion/reabsorption pathways were identified as most significantly enriched by this analysis. The size of the circle adjacent to each pathway indicates the number of differentially expressed genes (DEG) within that pathway; and the color indicates the -$\text{Log}_{10}$ of the p-value for the enrichment. **b** and **d**, Violin plots comparing the level of expression of mRNAs between cluster 0 and cluster 3 cells that are components of the JAK-STAT signaling pathway (*LIFR, CNTFR, PDGFRB, STAT3, SOCS1, SOCS3, MCL1* and *PIM1*)**,** TP53[70] and cellular senescence (*GADD45B*[71], *GADD45G, IGFBP3, SERPINE1, THBS1, TGFB1, TGFBI, SQSTM1, NFKB1, CXCL8* and *EIF4EBP1*), and in the regulation of stem cell pluripotency (*LIFR, STAT3, INHBA, ID1, ID2, ID3, FGFR1* and *PCGF2*). **c**, Violin plots comparing the level of expression of *LIF, PDGFA* and *PDGFB* mRNAs in all 15 clusters in day 21 organoids. **e**, Violin plot showing the level of *STAT3* mRNA expression during HO development. scRNA-Seq data was generated from iPSC, day 9 hepatoblast (HB), primary (HO1) and secondary organoids (HO2) cultured in growth media (GM) or in differentiation media (DM). Primary human hepatocyte (PHH) mRNA was used as control.

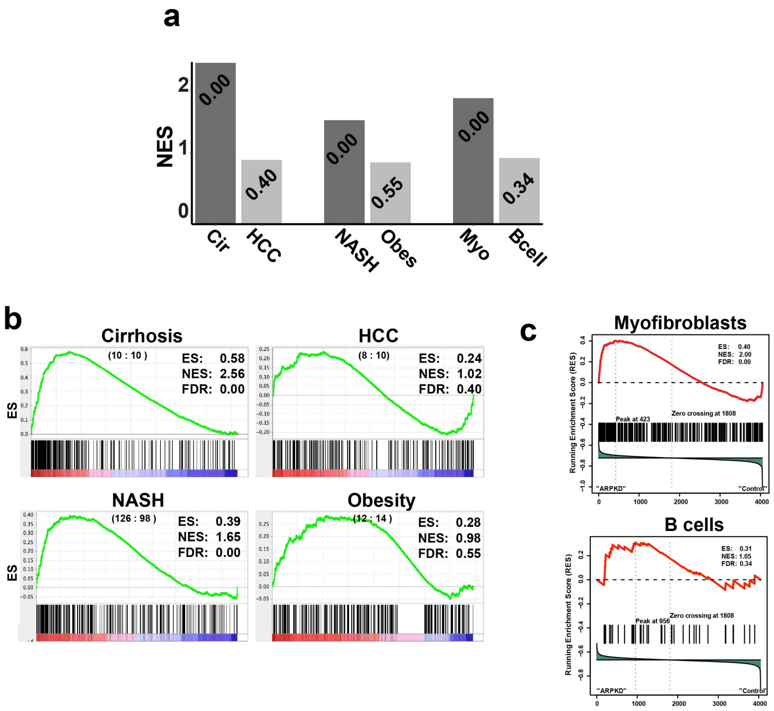


**Figure S9**. **ARPKD organoid myofibroblasts resemble those present in human liver tissue with commonly occurring forms of human liver fibrosis.** The ARPKD organoid myofibroblast gene expression signature is present in liver tissue obtained from patients with commonly occurring forms of liver fibrosis. Gene signature expression analysis (GSEA) was performed to correlate the ARPKD organoid myofibroblast gene expression signature with that in liver tissue obtained from patients with cirrhosis (**Cir**), nonalcoholic steatohepatitis (**NASH)-** induced fibrosis, hepatocellular carcinomas without fibrosis (**HCC**), or from obese subjects (**Obes**) without fibrosis. **a,** The bar graph shows the GSEA results (normalized enrichment score (NES)) obtained from each analysis; and the false discovery rate (FDR) for each comparison is shown at the top of each bar. The myofibroblast signature was very strongly associated with cirrhotic and NASH liver tissue, but it was not induced by obesity alone or by the presence of a HCC. GSEA was also performed by comparing the cluster 0 cell transcriptome with overlapping genes present in the expression signatures defined from the scRNA-Seq analysis of myofibroblasts and B cells present in cirrhotic human livers [24]. **b**, GSEA results for the correlation of the myofibroblast gene expression signature in ARPKD organoids with that in liver tissues obtained from: 10 normal and 10 cirrhotic subjects (**Cirrhosis**); 98 normal and 126 NASH subjects (**NASH**); 10 normal and 8 hepatocellular carcinomas (**HCC**); and 14 subjects with normal body weight and 12 obese subjects (**Obesity**). The myofibroblast signature was very strongly associated with cirrhotic and NASH liver tissue, but it was not induced by obesity alone or by the presence of a HCC. On the x-axis: each vertical line represents one of the 254 genes in the myofibroblast expression signature; a red color at the bottom of the graph indicates the expression for that gene is positively correlated with expression in the indicated disease tissue, while a blue color indicates correlation with the normal condition. The Enrichment Score (ES, y-axis) indicates the correlation of each gene with the indicated disease condition. The values for the ES, the normalized enrichment score (NES), and the false discovery rate (FDR) for each of the four comparisons are shown. **c**, As controls, GSEA was also performed by comparing the genes present in cluster 0 cells with overlapping genes in the expression signatures of myofibroblasts or B cells; which were identified by scRNA-Seq analysis of cirrhotic human livers [24]. The myofibroblast gene signature found in cirrhotic human liver tissue was strongly correlated with ARPKD organoids but not with control organoids. As additional controls, GSEA results indicated that the B cell signature was not abundant in either control or ARPKD organoids; and that the hepatocyte and cholangiocyte expression signatures were evenly distributed between control and ARPKD organoids.

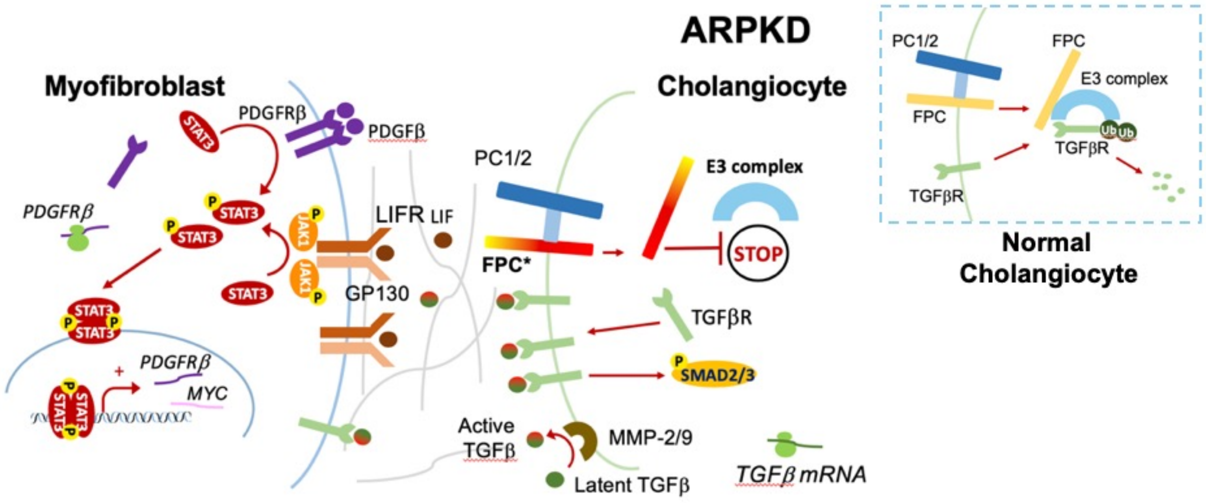


**Figure S10**. A proposed model for the pathogenesis of ARPKD liver disease. Within a normal cholangiocyte (dotted box), the FPC-mediated interaction between the TGFβ receptor and the E3-ubiquitin complex will lead to TGFβ receptor (TGFβR) degradation. The ARPKD mutation (FPC*) interferes with this interaction, which inhibits TGFβR degradation in ARPKD cholangiocytes. ARPKD cholangiocytes also have an increased level of MMP-2 and MMP-9 expression, which converts latent TGFβ to its active form. Increased TGFβ and TGFβR expression activates TGFβ-associated signaling pathways in ARPKD cholangiocytes. TGFβ acts in concert with LIF and PDGFβ, which are constitutively produced by multiple different cells types, to promote mesenchymal cell differentiation into collagen-producing myofibroblasts. The myofibroblasts, which have increased levels of LIF receptor and PDGFRβ receptor expression and STAT3 pathway activation, mediate the pathogenesis of ARPKD liver fibrosis.
